# Canonical decision computations underlie behavioral and neural signatures of cooperation in primates

**DOI:** 10.1101/2025.10.22.683999

**Authors:** Weikang Shi, Olivia C. Meisner, Monika P. Jadi, Steve W. C. Chang, Anirvan S. Nandy

## Abstract

Successful cooperation requires dynamic integration of social cues. However, the neural mechanisms supporting this complex process remain unknown. Here, we reveal that the primate dorsomedial prefrontal cortex (dmPFC) implements a gaze-dependent social evidence accumulation process to guide cooperative decisions in freely moving marmoset dyads. A drift-diffusion process in which the partner’s action variability is accumulated by social gaze best explains the cooperative actions of the actor. Single-neuron recordings in dmPFC revealed a direct neural correlate: the slope of predictive ramping activity mapped directly onto the rate of evidence accumulation, while baseline firing, modulated by prior outcomes, mapped onto the initial bias. At the population level, the geometry of dmPFC neural trajectories reflected the strength of social evidence and was linked to cooperative success. Together, these findings establish a multi-level neural mechanism for transforming active sensing into a decision variable, linking a canonical computation to cooperative behavior in a naturalistic setting.

## Introduction

Cooperation is a defining feature of advanced social behavior. To cooperate successfully, individuals must monitor the actions of their partners, infer intentions, and decide whether to act in a coordinated manner ^1^. A central challenge lies in transforming social cues—particularly through gaze, which reveals attention and intention ^2,3^—into decision variables that shape action selection.

Evidence accumulation frameworks provide a powerful explanation for how the brain integrates information to form a decision ^4,5^. Such mechanisms are well-established in domains such as value-based choice ^6–8^ and, most notably, perceptual decision-making ^5,9^. In perceptual tasks, neural correlates of evidence accumulation have been identified in regions like the dorsolateral prefrontal cortex (dlPFC) ^10^ and the lateral intraparietal area (LIP) ^11^, where the neural activity encodes the strength of incoming sensory evidence. However, two fundamental challenges remain. First, it is unclear whether evidence accumulation is a core, conserved computational mechanism that extends to the domain of complex social cognition. Second, because prior studies have relied on highly constrained, head-restrained task structures, it is unknown if these accumulation processes operate during naturalistic, unconstrained social interactions.

To address these challenges, we designed a cooperation task where freely moving pairs of marmosets coordinate their actions for mutual reward by pulling their lever within a 1-second cooperative time window (**Fig. 1A**) ^12^, allowing us to investigate the process of social decision-making in a naturalistic setting. While our recent work has identified the use of gaze-dependent strategies during these cooperative interactions ^13^, the precise manner in which gaze is used to integrate available social information and the neural mechanisms underlying this integration process remain unknown. In particular, although social gaze was necessary for successful cooperation, the cooperative time window was too temporally constrained to support a simple strategy in which the actor observes the partner’s pull and then reactively initiates its own pull ^13^. We therefore hypothesized that during these naturalistic social interactions, social gaze allows the actor to sample behaviorally relevant information from the partner’s ongoing actions, information that is predictive of the partner’s forthcoming cooperative behavior, and to integrate this information over time to guide the timing of its own action. Consistent with this view, we further hypothesized that during these naturalistic social interactions, the animals self-organize their behavior into discrete, self-paced episodes (henceforth “trials”) ^14^, each consisting of a period of gaze-dependent social evidence accumulation culminating in a cooperative decision. Conceptually, we defined each trial as a complete behavioral epoch, beginning with the initiation of movement leading to a pull action and ending after the outcome of that action was delivered and evaluated. To investigate the neural underpinnings of this process, we recorded from the dorsomedial prefrontal cortex (dmPFC), a key node in the social brain network ^15–17^. Given the dmPFC’s involvement in encoding others’ actions and outcomes ^18^, we hypothesized that it plays a key role in the accumulation of social evidence. By combining behavioral modeling with single-neuron and population-level analyses, we provide the first evidence demonstrating that the dmPFC implements a social gaze-dependent evidence accumulation process to transform social information into decision variables guiding cooperation (**Fig. 1A**).

**Figure 1.**
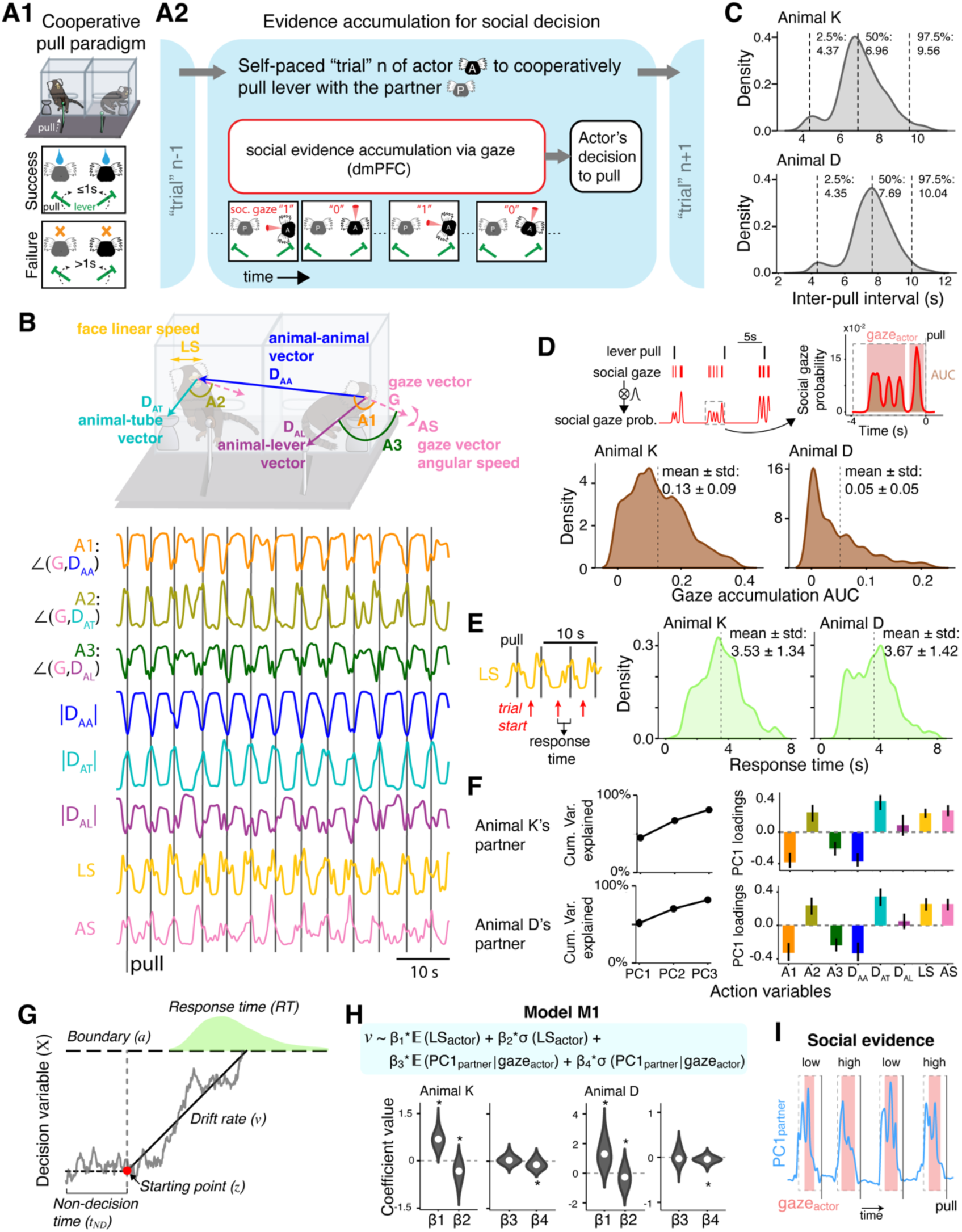
Conceptual hypothesis, behavioral metrics, and drift diffusion model. (**A**) Task design and the conceptual hypothesis: (**A1**) In the mutual cooperation (MC) task, marmoset dyads are required to pull their levers within a 1-second time window to receive a juice reward for both. (**A2**) We hypothesize that the social gaze of the actor animal collects social evidence about a partner’s cooperative intention, which is accumulated by the dorsomedial prefrontal cortex to guide action decisions. (**B**) Top: Definition of eight continuous behavioral variables. Bottom: Example traces for one animal in one session. (**C**) Distribution of inter-pull intervals. (**D**) Definition of the social gaze accumulation as the area under curve (AUC) of the social gaze probability. Dashed rectangle shows 4 s functional window of analysis. Pink shaded areas highlight social gaze duration (social gaze probability> 0). (**E**) Definition of the trial start and response time (RT). (**F**) Definition of the holistic representation of the partner’s actions via principal component analysis (PCA). (**G**) Structure of the drift diffusion model. (**H**) Coefficient estimates from the main model (M1) showing that partner action (PC1) contributes only when social gaze is present. (**I**) Definition of the social evidence. Wilcoxon signed-rank test for panel H (* p < 0.05). See also Supplementary Figures S1-S4, Movie S1 and Table S1.

## Results

### A quantitative framework for studying naturalistic cooperation

To quantify the animals’ behavior in the mutual cooperation task, we first used markerless facial tracking to extract eight continuous behavior variables for each marmoset ^19,20^. These variables captured three broad aspects of task-relevant behavior: (i) spatial relationships between the animal’s face and key task elements (the partner, the lever, and the juice tube), (ii) gaze orientation, defined by the angle between the head-gaze vector and these task-relevant targets, and (iii) movement dynamics, including linear and angular speed of the face (**Fig. 1B; Supp. Mov. S1**). We emphasize that this set of variables was not intended to exhaustively describe body posture or limb-level movements, but to capture task-relevant spatial relationships, gaze orientation, and movement dynamics that precede cooperative actions in a functionally meaningful time frame. Analysis of the intervals between consecutive pulls by each animal revealed a rhythmic pattern, supporting the characterization of the interactions as a series of self-paced trials (**Fig. 1C**). For both animals from which we recorded the neural activity (hereafter, the ‘actors’), the majority of these inter-pull intervals were longer than 4 seconds, a timescale that informed the functional window for our subsequent analyses. Within each trial, we defined “social gaze” as the actor looking at the partner’s face, and social gaze accumulation as the area under the curve of the social gaze probability in this functional window preceding a pull action (**Fig. 1D**). This measure revealed substantial variability in the amount of social gaze expressed before the pulls.

Inspection of the average time course of all eight behavioral variables aligned to pull onset revealed a gradual ramping of the linear speed (LS) beginning several seconds before the pull (**Supp. Fig. S1A**). Other variables also exhibited systematic average changes consistent with approaching the lever and partner and reorienting gaze. Despite these average tendencies, there was substantial variability across pulls: median traces had large interquartile ranges, and pairwise correlations across single-pull traces spanned nearly the full range from −1 to 1 (**Supp. Fig. S1A**). To compare the relative timing of these behavioral changes, we analyzed rescaled median traces and deviation times on a pull-by-pull basis (**Supp. Fig. S1B–C**). Although LS tended to ramp earlier on average, deviation times were broadly distributed and highly overlapping across variables, indicating that no variable exhibited a narrow or invariant earliest deviation time. Thus, pre-pull behavior does not follow a stereotyped or strictly ordered temporal sequence. To objectively define the start of each trial, we therefore sought the single most powerful and unique behavioral predictor of a future pull. Using a Generalized Linear Model (GLM), we found that the linear speed (LS) of the actor’s face was a consistently high-importance predictor in both animals (**Supp. Fig. S2A**). Furthermore, a Variance Inflation Factor (VIF) analysis confirmed that LS had low collinearity with the other variables, indicating it provided unique predictive information (**Supp. Fig. S2B**). Based on this evidence, we used the ramping onset of face linear speed to define the start of each trial, and operationalized the duration from this starting point to the pull action as the response time (RT) for that trial (**Fig. 1E**). To further characterize within-trial structure, we examined how social gaze was distributed across the trial. Rather than occurring continuously, the gaze probability signal was segmented into multiple discrete bouts with variable timing and duration relative to pull onset (**Table S1**). Across trials, animals showed substantial variability in the number and duration of these bouts, as well as in the proportion of time spent gazing at the partner. This heterogeneity indicates that cooperative behavior cannot be captured by a single gaze event or by a fixed gaze-to-pull interval.

With this quantitative framework in place, we hypothesized that the actor animal utilizes a holistic representation of the partner’s actions that would collectively allow it to infer the partner’s cooperative intention. This hypothesis is motivated in part by the substantial trial-to-trial heterogeneity observed during pre-pull behavior (**Supp. Fig. S1**), suggesting that the decision process must flexibly rely on context-dependent cues rather than be governed by a fixed sequence of events. To define such a representation, we performed Principal Component Analysis (PCA) on the partner’s eight behavioral variables. PCA was performed separately for each behavioral session to avoid pooling across potentially heterogeneous interaction dynamics. The first principal component (PC1) captured a large fraction of the behavioral variance and thus served as a summary of the partner’s actions (**Fig. 1F**). Importantly, PC1 does not encode a specific behavioral event or state; rather, it provides a low-dimensional summary of dominant fluctuations in partner behavior at each moment, allowing diverse and heterogeneous behavioral cues to be represented within a common framework. PC1 showed relatively uniform loadings across nearly all eight behavioral dimensions, with no single variable dominating the component, indicating that PC1 summarizes global partner activity rather than reflecting a specific feature. Importantly, this loading structure was highly similar across partners and sessions (**Fig. 1F**). This pipeline thus allowed us to deconstruct continuous behavior into a structured, trial-based format suitable for studying the social evidence accumulation process.

### Gaze-dependent drift-diffusion model explains cooperative decisions

To formalize the decision-making process underlying cooperation, we fit the animals’ response times and choices using a drift-diffusion model (DDM) ^21^, which decomposes behavior into several key parameters: drift rate (*v*, the speed of accumulation), the starting point (*z*, the initial bias), and non-decision time (*t_ND_*) (**Fig. 1G; see Methods**). Critically, we formulated four competing models to explicitly test how social gaze mediates the integration of partner information (**Supp. Fig. S3A**). Model comparison using the Deviance Information Criterion (DIC) revealed that a model which allows partner evidence to be integrated only during period of social gaze (M1; **Fig. 1H**) provided the best fit compared to alternate models assuming either no gaze modulation (M2), or an additive (M3), or multiplicative (M4) effect of gaze accumulation (**Supp. Fig. S3B**). Importantly, the resulting accumulation dynamics were qualitatively consistent across pairings with different partners, with the direction of the social evidence effect on drift rate remaining negative across all tested pairs (**Supp. Fig. S3C**). This result supports a gaze-dependent integration mechanism, where information about the partner’s actions is specifically sampled during periods of social attention.

Analysis of the coefficients from the best-fitting model (M1) provided crucial insight into the nature of the social evidence being accumulated. We found that the coefficient for the variability of the partner’s actions (*β*_4_) was consistently and significantly negative in both animals (**Fig. 1H**). Coefficients capturing the actor’s own action variables (*β*_1_ and *β*_2_) were also significant, consistent with the fact that the model predicts the actor’s pull behavior, whereas the coefficient capturing the mean level of the partner’s actions (*β*_3_) was not. Importantly, the effect of *β*_4_ remained significant after accounting for these other variables. This indicates that the variability of the partner’s action (PC1) was the key social information being accumulated, where a noisier or more ambiguous action (higher standard deviation) leads to a smaller drift rate, slowing the decision process. More specifically, in this cooperative lever-pulling task, the mean level of a partner’s action provides limited information about when the partner is likely to initiate a pull, whereas reduced variability reflects greater temporal regularity and predictability of the partner’s behavior, which directly supports coordinated action. Replacing the holistic PC1 with the partner’s linear speed—the only single variable identified to be both informative and non-redundant in our GLM and collinearity analyses—did not improve model performance, and its key coefficients were not significant (**Supp. Fig. S3D-E**). These results demonstrate that the decision process relies on a rich, multidimensional representation of partner behavior, rather than on any single feature.

These analyses suggest that, within each trial, social gaze tracks the moment-by-moment variability in the partner’s ongoing actions. However, actors could in principle rely on simpler cues, such as systematic trends in the partner’s actions or detecting a particular coincidence or threshold state, that might also produce the observed negative *β_4_* effect. To directly test such alternative possibilities, we examined whether partner behavior during the actor’s pre-pull social gaze exhibited consistent monotonic trends or converged to fixed terminal states that could serve as coincidence or threshold cues. As shown in Supplementary Figure S4, neither the slopes of partner behavioral variables during social gaze (**Supp. Fig. S4A**) nor their values at the end of gaze (**Supp. Fig. S4B**) showed consistent structure across trials, ruling out models based on trend detection or fixed state thresholds. Accordingly, we define “social evidence” in each trial as the within-trial variability (standard deviation) of the partner’s PC1 during periods of social gaze, such that lower variability (less noise) corresponds to stronger social evidence (**Fig. 1I**; see **Methods**). In this framework, accumulation refers to the integration of uncertain social information over time to guide the actor’s decision, rather than to coincidence or state detection.

### dmPFC neurons show predictive, mixed selectivity for social variables

To investigate the neural basis of this social gaze-dependent process, we recorded single-unit activity from the dmPFC of the actor marmosets using a custom-built, wireless recording system (**Fig. 2A; Supp. Mov. S1**). We implemented a GLM (**Fig. 2B; see Methods**) to formally characterize the diversity of observed neural responses (**Supp. Fig. S5A**). Many dmPFC neurons exhibited mixed selectivity, including a core subpopulation of “triple-encoding” neurons that simultaneously represented all three key task variables: the actor’s pull, the actor’s gaze, and the partner’s actions (**Fig. 2C**).

**Figure 2.**
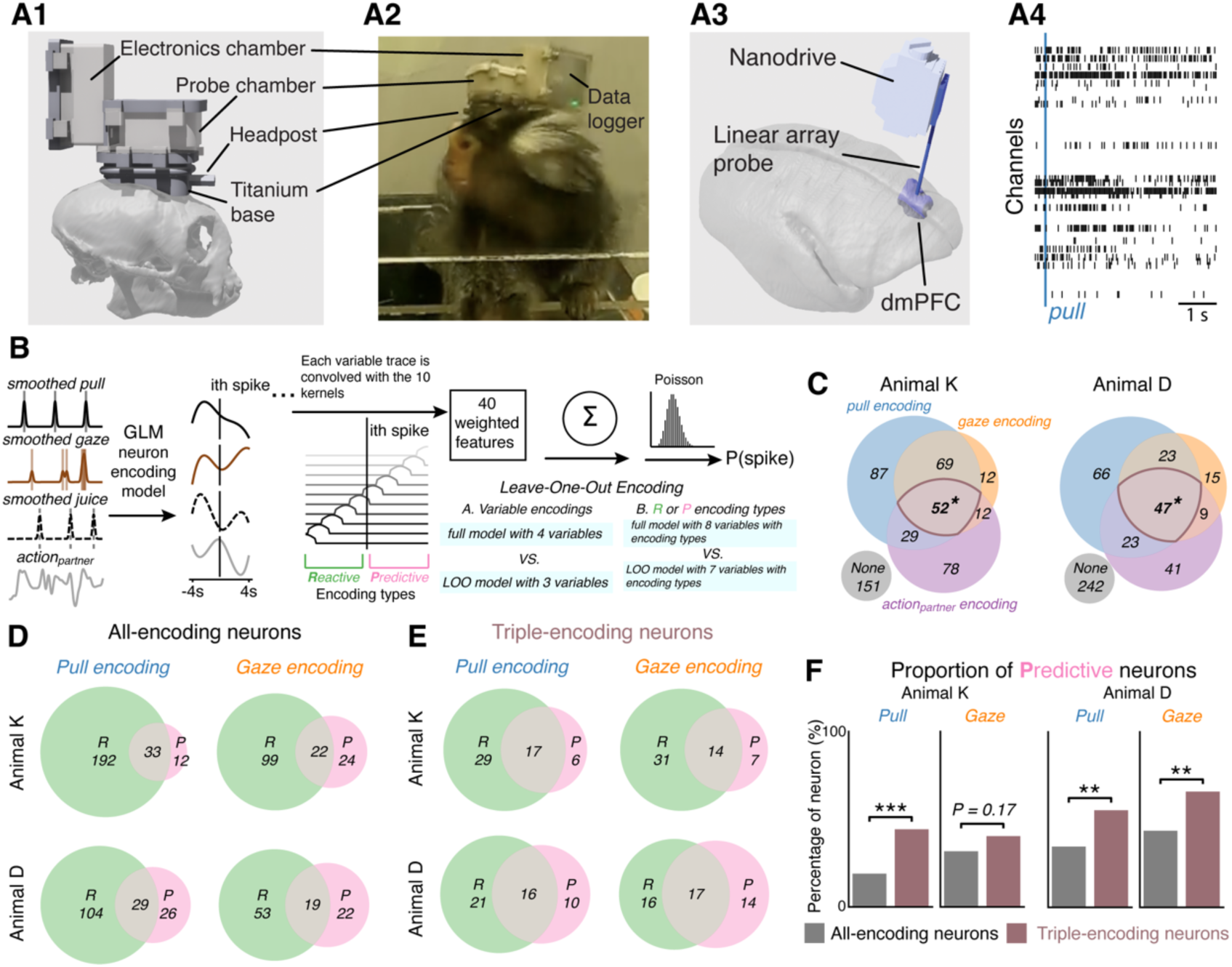
dmPFC neurons show mixed selectivity. dmPFC neurons integrate self, partner, pull and gaze variables, and encode both predictive and reactive components of the actions. (**A**) Wireless recording system design: (**A1**) illustration of the headcap design; (**A2**) example video frame highlighting the wireless recording apparatus on one of the animals during task performance; (**A3**) estimation of the dmPFC recording sites; (**A4**) example of action potentials (black ticks) from the 64-channel probe aligned at a pull action. (**B**) GLM pipeline for defining tuning properties and encoding types. (**C**) Distribution of selectivity across neurons reveals mixed coding for pull, gaze, and partner action variables, supporting flexible social computations. (**D**) Distribution of encoding categories for pull and gaze neurons. R, reactive; P, predictive. (**E**) Subset of “triple-encoding” neurons’ coding categories for pull and gaze actions. (**F**) Predictive encoding is enriched among triple-encoding neurons. Chi-squared test for panel C and Fisher’s exact test for panel F (* p < 0.05, ** p <0.01, *** p< 0.001). See also Supplementary Figure S5.

We next investigated the temporal dynamics of this neural encoding, refining our GLM to classify tuning as either “predictive” (neural spikes preceding behaviors) or “reactive” (neural spikes reacting to behaviors) (**Fig. 2B; see Methods**). We found distinct subpopulations with purely predictive or reactive profiles, as well as many that exhibited both, consistent with mixed selectivity in dmPFC (**Fig. 2D-E; Supp. Fig. S5B**). Importantly, this classification is based on multivariate statistical relationships rather than requiring neural activity to occur exclusively before or after a given event at the level of individual neurons, which is particularly relevant in this task, where gaze, pull, and outcome variables are temporally correlated. Consistent with this interpretation, examination of single-unit activity and population distributions (**Supp. Fig. S5B-C**) revealed heterogeneity at the level of individual neurons, but a clear population-level structure: neurons classified as predictive exhibited earlier activity relative to the aligned event, whereas reactive neurons showed delayed modulation. Critically, we reasoned that neurons encoding the most comprehensive set of task variables would be particularly involved in linking sampled social information to the actor’s own actions. Supporting this idea, the triple-encoding subpopulation was significantly enriched for predictive encoding of pull and gaze actions relative to the broader population of task-responsive neurons (**Fig. 2F**). Together, these findings indicate that dmPFC neurons integrate social information and self-actions in a statistically anticipatory manner, with population-level predictive signals emerging prior to cooperative actions rather than solely reflecting reactive responses.

### Neural ramping activity reflects evidence accumulation

Having established that dmPFC neurons show predictive, mixed selectivity, we next asked whether their activity exhibits the dynamic signatures of the evidence accumulation process itself. A key feature of a neural accumulator is ramping activity, wherein the slope is inversely related to RT. We found precisely this relationship in the dmPFC population: the average slope of the pre-pull firing rate was steeper for faster responses and shallower for slower ones (**Fig. 3A**). Critically, these firing rate slopes scaled with the strength of social evidence (**Fig. 3B**). A significant subpopulation of neurons conjunctively encoded both RT and social evidence in their firing rate slopes (**Fig. 3C**). Crucially, this encoding of social evidence by differential ramping was strongly linked to behavioral outcome: this relationship was robust on successful trials but significantly weaker on failed trials (**Fig. 3D**). Notably, the magnitude of the ramping slope itself did not predict choice correctness once response time was controlled (**Supp. Fig. S6**), indicating that dmPFC ramping does not directly encode success or failure. Instead, behavioral outcome depended on the degree to which ramping dynamics were aligned with social evidence, consistent with an accumulation process that guides action timing rather than explicitly signaling correctness. We further found a strong correspondence between this dynamic slope-based encoding and the static tuning properties of the neurons (**Supp. Fig. S7A**). Moreover, this slope-encoding population showed a trend of being enriched for predictive tuning, particularly for pull-predictive neurons, which is consistent with an anticipatory, rather than reactive, process (**Supp. Fig. S7B**). Consistent with this anticipatory interpretation, peak firing times of individual dmPFC neurons were typically close to the lever pull (**Supp. Fig. S8A**) and relatively stable across trials (**Supp. Fig. S8B**), suggesting that the slope effects in Figure 3A-D are less likely to reflect temporally misaligned step-like responses.

**Figure 3.**
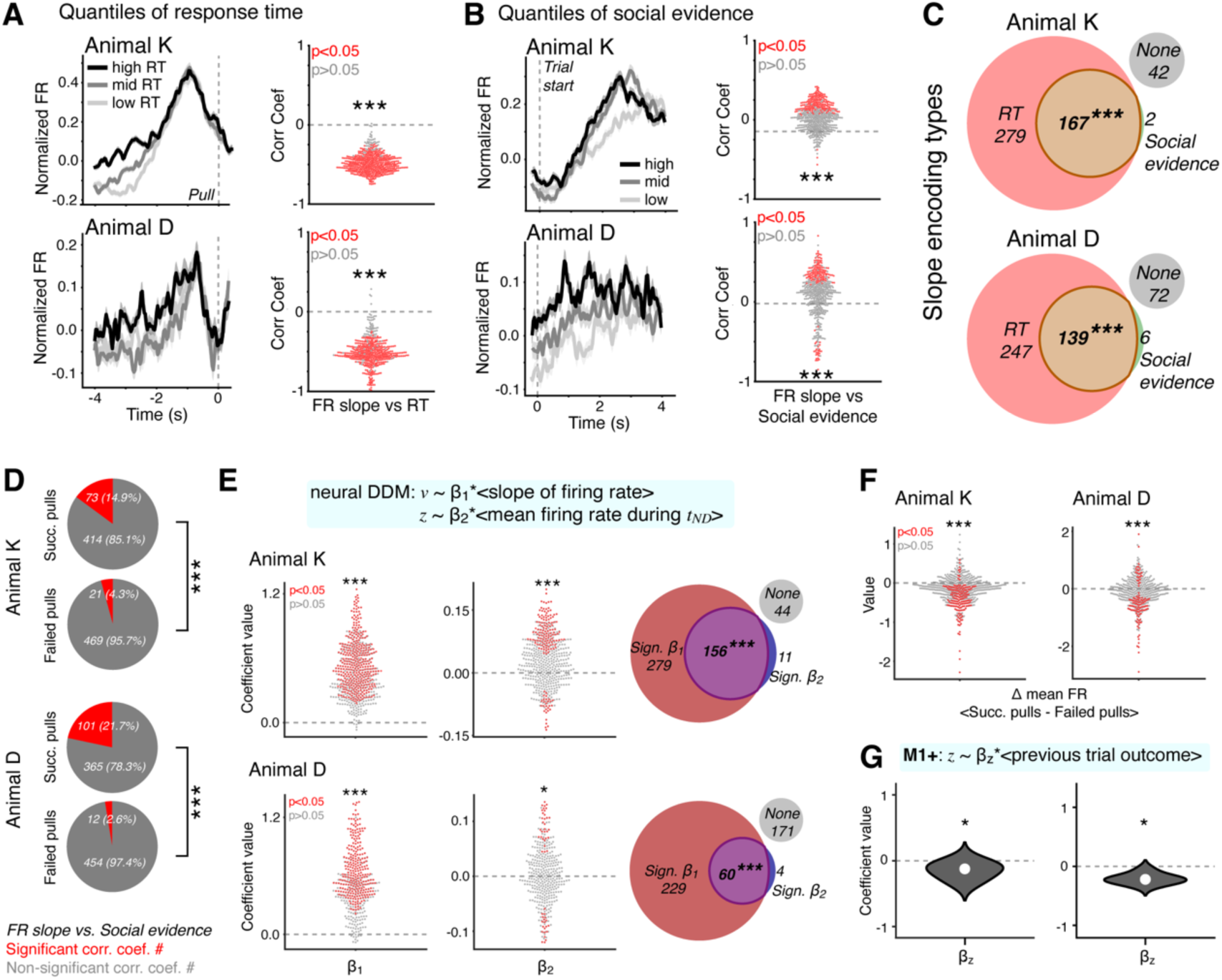
Neural firing rate slopes encode both response time and social evidence. (**A**) Mean normalized firing rate of all dmPFC neurons aligned at the pull action and separating into three quantiles of the RTs (left), and correlations between dmPFC firing rate slopes and RT (right). (**B**) Mean normalized firing rate of dmPFC neurons aligned at the trial start and separating into three quantiles of the social evidence (left), and correlations between dmPFC firing rate slopes and RT social evidence (right). (**C**) Significant proportions of neurons encode both factors simultaneously. (**D**) Neuron numbers with and without significant correlations between firing rate slopes and social evidence in successful vs failed pulls. € Neural DDM linking firing rate slope to drift rate (*v*) and baseline firing to starting point (*z*). (**F**) The difference in the mean baseline firing rates following successful pulls vs. failed pulls. (**G**) Behavioral DDM model incorporating outcome-dependent starting points. Wilcoxon signed-rank test for panels A, B, E, F and G, Chi-squared test for panels C and E, and McNemar test for panel D (* p < 0.05, ** p <0.01, *** p< 0.001). For the red datapoints, Pearson’s correlation test for panels A and B, 95% Credible Interval (CI) of the posterior distributions for panel E, and Mann-Whitney U Test for panel F. See also Supplementary Figure S6-S8.

### Neural activity maps directly onto computational model parameters

To formally connect the observed neural dynamics to the computational model, we constructed a neural DDM that incorporated both the firing rate slopes and baseline firing rates as predictors. The results revealed a direct mapping: trial-by-trial firing rate slopes were a significant positive predictor of the DDM’s drift rate, while baseline firing rates quantified during the non-decision time predicted the starting point (**Fig. 3E**). This starting point bias was itself influenced by recent history, in which a failed prior trial was associated with higher baseline firing (**Fig. 3F**). To test the hypothesis suggested by this neural result—that post-error changes in baseline activity have a direct behavioral consequence—we returned to our best behavioral model (M1) and added the previous trial’s outcome (1 for successful pulls and 0 for failed pull) as a predictor of the starting point. This updated model (M1+) confirmed the hypothesis, revealing a significant negative coefficient for this relationship (**Fig. 3G**). This finding provides a direct behavioral correlate for the neural activity, showing that the outcome of the prior trial systematically biased the starting point of the current decision process, potentially reflecting a post-error increase in motivation. These analyses provide a direct bridge between the firing rate and computational levels, demonstrating that the dynamic and static components of dmPFC activity map onto the core parameters of the evidence accumulation process.

### Population dynamics link neural processing to cooperative success

While single-neuron activity correlates with key decision variables, cognitive functions are ultimately implemented by the coordinated, dynamic activity of large neural populations. Therefore, to understand how the dmPFC as a whole processes social evidence, we examined the collective neural activity by analyzing population trajectories at the level of single trials (**Fig. 4A**). We found that the geometry of these neural population trajectories, specifically their path variability, defined as the local tortuosity of the trajectory ^22^, was significantly negatively correlated with the level of social evidence (**Fig. 4B-C**). This relationship was most pronounced early in the trial (**Fig. 4B**), a period during which social information is actively sampled, although we note that the overall temporal profile of path variability across the trial (**Fig. 4E**) is descriptive and may reflect generic early-to-late convergence of neural activity as actions are approached. Crucially, the evidence–path variability relationship was present only on successful trials and was absent on failed trials (**Fig. 4D**), despite similar gross motor actions occurring in both cases. At the population level, this outcome dependence mirrors our single-neuron results, where social-evidence encoding in firing-rate slopes was robust on successful trials but significantly weaker on failed trials (**Fig. 3D**). Together, these population-level findings offer converging evidence that dmPFC dynamically encodes social evidence in a manner that supports successful cooperation.

**Figure 4.**
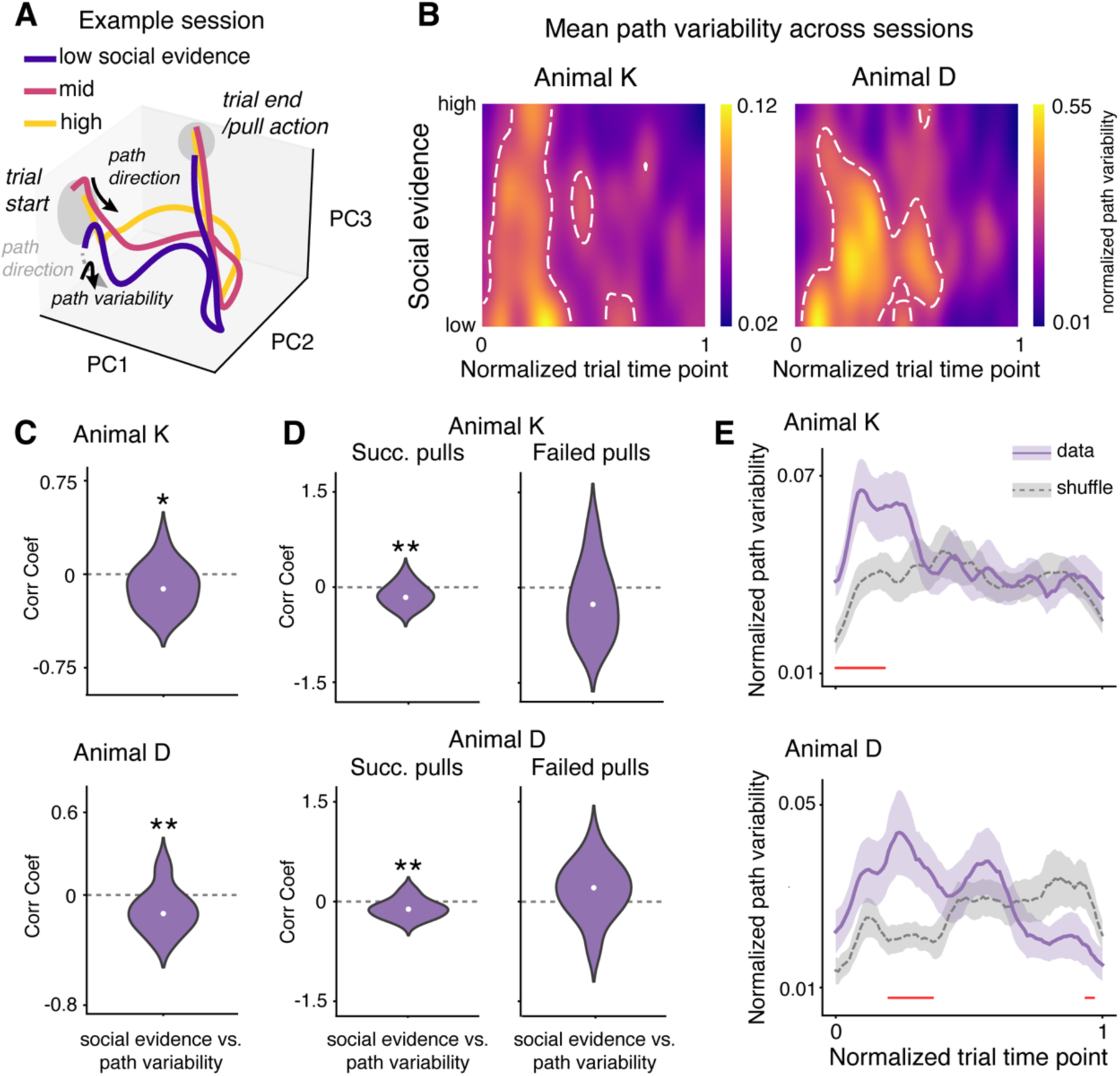
Population dynamics reflect social gaze accumulation. (**A**) Definition of path variability (tortuosity) of the single-trial neural population trajectory, example session. (**B**) Heatmaps of the temporal structure of the path variability showing that social evidence shapes dmPFC population dynamics during social decisions, averaged across sessions. (**C**) Correlation between the strength of social evidence and path variability of the neural trajectory across sessions. (**D**) Correlation between social evidence and path variability of the neural trajectory in successful and failed pulls. € Average change of path variability over time in individual trials. Wilcoxon signed-rank test for panels C and D (* p < 0.05, ** p <0.01), and paired t-test with cluster-based correction for panel E (red segments p < 0.05).

## Discussion

Our study provides a multi-level account of how the primate dmPFC orchestrates cooperation in a naturalistic behavioral setting. We show that the dmPFC implements a social gaze-dependent evidence accumulation process, transforming actively sampled social cues into a decision variable. By identifying this canonical computation in freely moving animals without any imposed task structure, our work bridges the gap between classic decision neuroscience and the complexity of real-world social interactions.

Our findings connect behavior, computation, and neural activity across multiple scales. A gaze-dependent DDM best explained cooperative decisions, establishing that evidence about a partner’s actions is integrated only when actively sampled. We then identified a direct neural correlate for this process. At the single-neuron level, the predictive ramping of dmPFC neurons encoded both the response time and the strength of social evidence. These neural dynamics mapped directly onto the DDM’s core parameters, with firing rate slopes predicting the drift rate and baseline activity—updated by recent trial outcomes—predicting the starting point bias. Finally, this mechanism was reflected at the population level, where the geometry of neural trajectories tracked the level of social evidence. Crucially, this signature was present mostly on successful trials, cementing the link between the neural mechanism and the behavioral outcome.

Evidence accumulation has long been established as a canonical computation for perceptual decisions ^5^, with clear neural signatures in sensorimotor areas such as LIP ^11^ and dlPFC ^10^. Our findings extend this framework into the social domain, identifying the dmPFC as a key locus for this more complex form of accumulation. Unlike perceptual tasks, where evidence is often externally driven, social evidence about another’s intentions is uncertain and must be actively sampled via gaze ^2^. In this context, social gaze appears to sample partner behavior that is heterogeneous and trial-dependent, making it difficult to rely on any single momentary signal. Instead, the relevant variable reflects the reliability of partner behavior within a trial—analogous to motion coherence in perceptual decision-making tasks, where coherence is fixed per trial and determines decision difficulty rather than serving as a time-varying feature to be detected. Consistent with this interpretation, our analyses rule out models based on monotonic trend detection or convergence to specific behavioral states, indicating that accumulation reflects the integration of uncertain social information over time rather than coincidence or threshold detection. The dmPFC is uniquely suited for this computation ^23^, integrating information about the partner’s actions with the actor’s own social attention. Importantly, we do not interpret these findings as implying that dmPFC uniquely implements social evidence accumulation; rather, dmPFC likely participates in a broader frontal network in which social information is integrated into action planning to guide behavior.

A significant advance of our study is the application of a detailed, mechanistic approach to neural function in freely moving, socially interacting primates. While the field has begun to embrace the study of more naturalistic behaviors ^24^, as exemplified by recent work in rodents ^25^, tree shrews ^26^, and macaques ^27^, research on the detailed computational mechanism has been lacking, facing both technical hurdles in recording from unconstrained animals and analytical challenges in interpreting complex, continuous behavior. We addressed the technical challenges by developing a novel modular head chamber ^28^, which improved surgical success rates. We also used movable probes, in contrast to fixed chronic implants ^29^, to sample diverse neural populations across sessions. To overcome the analytical challenges, we employed a sophisticated computational framework, including the GLM pipelines ^30–33^, to deconstruct the continuous behavioral stream into meaningful, quantifiable variables. Importantly, applying this framework revealed a distinction within dmPFC between neurons that reactively track ongoing behavioral and social variables and neurons whose activity predicts upcoming self-actions while also encoding partner-related information. This distinction helps dissociate anticipatory social computation from simple movement planning signals and clarifies dmPFC’s role in linking sampled social information to forthcoming decisions. By identifying computational signatures of decision-making in this unconstrained paradigm, we show that even without the rigid structure of a classic task, marmosets spontaneously organize their behavior in a way that is amenable to an evidence accumulation process. This demonstrates the robustness of this neural algorithm, suggesting it is a fundamental process that is flexibly deployed to make decisions in dynamic environments.

Although the present study focuses on a specific cooperative pulling task, the experimental and analytical framework is intended to capture a more general process in social behavior. In many social contexts, individuals must continuously accumulate social information and align it with the timing and preparation of their own actions, as in observational learning ^34^ or collective decisions ^35^. From this perspective, we suggest that dynamic alignment between social evidence and action preparation represents a general motif through which medial frontal circuits support flexible social coordination across tasks and species.

Our results also highlight several avenues for future research. The influence of the previous trial’s outcome on the DDM starting point, mediated by baseline dmPFC firing, suggests a rapid, experience-dependent plasticity that could be interpreted as an alerting signal or a shift in motivation following failure ^36–38^. Future studies employing neural manipulation could also directly test the causal role of this mechanism; for instance, microstimulation delivered during the gaze accumulation period could potentially disrupt the evidence integration process and alter the cooperative decision ^39^. In our modeling framework, social gaze serves as a gating signal that permits continuous accumulation of social evidence while gaze is present. Although this formulation captures the time-resolved influence of gaze on social evidence accumulation, it does not exclude the possibility that, at finer temporal scales, social information may exert its influence in a more threshold-like or non-linear manner within gaze bouts, an important question for future experimental and theoretical work.

In conclusion, by linking behavior, modeling, and neural dynamics across multiple scales, we established a comprehensive framework for how the brain transforms active sensing into a decision variable to guide cooperation. This neurocomputational framework provides a mechanistic grounding for economic models of strategic interaction ^40^, a potential biomarker for social deficits in psychiatry ^41^, and offers new biological insights for designing the next generation of cooperative multi-agent artificial systems ^42^.

## STAR Methods

### EXPERIMENTAL MODEL AND SUBJECT DETAILS

#### Animal subjects

Two common marmosets (*Callithrix jacchus*) participated in the study: a 7-year-old female (Animal K) and a 6.5-year-old male (Animal D). Animal K participated in 21 sessions with four different partners (2 females and 2 males), and Animal D participated in 29 sessions with three different partners (3 females). A detailed comparison of behavior and neural activity across different partners is beyond the scope of this study. However, to confirm that our findings were not confounded by partner identity, we performed alternative analyses using linear mixed-effects models, which included partner identity as a random effect. These confirmatory models did not change the statistical conclusions of the primary analyses reported throughout the manuscript. The animals were pair-housed, maintained on a 12-hour light-dark cycle, and food and water were removed from their home cages for 1–3 hours prior to testing. All experimental procedures were approved by the Yale Institutional Animal Care and Use Committee and complied with the National Institutes of Health Guide for the Care and Use of Laboratory Animals.

### METHOD DETAILS

#### Experimental task

The experimental apparatus and training procedures were identical to those described in previous studies ^12,13^. The animals had been previously trained to perform a mutual cooperation (MC) task. Both marmosets in the dyads were required to pull their respective levers within a 1-second time window to successfully receive a reward. A pull was registered when the lever passed a predefined position threshold, and its timestamp was recorded by the task-control computer at a 10 kHz sampling rate. After passing the threshold, the lever must be released (i.e., animals do not hold on to the lever) for it to be counted as a valid pull. A successful, coordinated trial resulted in the delivery of a 0.2 mL liquid reward (marshmallow fluff solution) to both animals, which was immediately preceded by an auditory tone (2 KHz) to signal success. A failed trial occurred if the pulls were not synchronized within the 1-second window. This task was designed to be performed in a naturalistic setting, with no explicitly defined or implemented temporal structure; instead, the animals engaged in a continuous series of self-paced cooperative decisions. The data analyzed in this study were collected after the animals had been fully trained on this task, representing established cooperative performance.

#### Behavioral Data Acquisition and Processing

Continuous behavior was recorded at 30 frames/sec from three synchronized cameras (GoPro Hero10) positioned at the front of the apparatus. The cameras were controlled via a shared remote control. To align the video data with the event timestamps from the task-control computer, we used two redundant signals, both controlled by the task computer: an audible tone at the start of the session for coarse alignment, and an LED that flashed every 20 seconds for fine alignment, providing a reliable and recurring synchronization signal captured by the cameras. We used the multi-animal version of DeepLabCut 2.0 to perform markerless pose estimation ^20,43^. Six facial key points were labeled on 500 frames: two ears, two eyes, the central blaze, and the mouth. The DeepLabCut neural network was trained on these labels for 150,000 iterations until the tracking error reached a stable minimum. The 2D coordinates from the three cameras were then used to triangulate the 3D position of each key point using Anipose ^19^. This 3D reconstruction allowed for the precise calculation of each animal’s head position and orientation in space throughout the session. From the 3D facial coordinates, we derived the eight continuous behavioral variables used for all subsequent analyses, chosen to capture task-relevant spatial relationships, gaze orientation, and movement dynamics rather than full-body posture (**Fig. 1B**):

- ***A1-A3***: Three angular variables measuring the direction of gaze relative to the other animal, the reward tube, and the lever.
- ***D_AA_, D_AT_, D_AL_***: Three distance variables measuring the distance to the other animal, the reward tube, and the lever.
- ***LS***: The face linear speed, calculated from the movement of the head’s 3D position.
- ***AS***: The gaze vector angular speed, capturing the rate of change in head orientation.

All spatial variables were derived from the 3D position of the face, operationalized as the centroid of the tracked facial landmarks, and therefore reflect face-based spatial relationships rather than full-body posture or limb position. We extracted these variables for each animal within a dyad, and they were computed separately for the actor and the partner. When defining the partner’s action and social evidence, we exclusively used variables derived from the partner’s behaviors; when defining the trial onset and response time, we exclusively used variables derived from the actor’s behaviors. Similar to earlier work ^13^, head gaze was defined as a 15-degree cone extending perpendicularly from the facial plane formed by the tracked key points. This definition reflects the limited range of eye-in-head movements in marmosets during naturalistic behavior and incorporates peripheral visual information ^44–46^. Social gaze was determined to have occurred whenever one animal’s gaze cone intersected with the other animal’s face. This frame-wise binary social gaze signal was sampled at the video frame rate (30 Hz) and smoothed using a Gaussian kernel (σ = 130 ms) to reduce frame-by-frame noise and yield a continuous social gaze probability signal (**Fig. 1D**). While our accumulation model operates on this continuous gaze signal as described below, we additionally quantified basic properties of discrete gaze bouts (defined as individual periods during which social gaze probability > 0) derived from the same signal for descriptive purposes. Summary statistics of gaze bout number, duration, and temporal distribution are reported in Supplementary Table S1.

#### Implant Design and Surgical Processing

We designed a novel, multi-component head implant to facilitate wireless neural recordings in freely moving marmosets (**Fig. 2A**). The system was custom-designed in CAD software (Shapr3D) to fit each animal’s cranium (obtained from pre-operative CT scan) and consists of two main parts. The first is a 3D-printed medical-grade titanium base (Materialise NV) that protects the surgical site where the skull is exposed. An integrated headpost on the base allows for brief periods of stable head-fixation. The second part is a 3D-printed recording chamber (nylon; Protolabs), which consists of two sealed and separated chambers. The probe chamber houses and protects a miniature screw microdrive (nanodrive; Cambridge Neurotechnologies) and the probe, while the electronics chamber protects the headstage chips needed for digitization and signal amplification. A silicone-based sealant (Kwik-Sil; World Precision Instruments, LLC) separates the chambers, allowing the probe to remain sterile while the data logger is connected to the headstage electronics. This modular design is advantageous for animal welfare, as it permits a multi-step surgical approach with shorter individual procedures and simplifies implant cleaning and maintenance.

Enabled by this modular implant design, the surgical implantation followed a multi-step process. First, skull geometry was extracted and reconstructed from the pre-operative CT scan to guide surgical planning and refine targeting. The 3D-printed titanium base was implanted, with its placement guided by the reconstructed skull and stereotaxic coordinates targeting the dorsomedial prefrontal cortex (dmPFC). In a subsequent procedure, a cranial window was opened within the protected base to access the dmPFC, with its location guided by the same CT reconstruction and stereotaxic coordinates. A nanodrive holding a 64-channel linear array probe (NeuroNexus) was mounted inside the probe chamber. The nanodrive allowed movement of the probe along the superior-inferior axis, while the mounting apparatus holding the nanodrive allowed repositioning along the anterior-posterior and mediolateral axes. The implant position was extracted from a post-operative CT scan. Although the CT resolution did not permit direct visualization of the probe shaft, the location of the nanodrive and planned trajectory were used to estimate probe position within dmPFC. The probe was lowered to a site within the dmPFC during the surgery, and the final recording site was confirmed post-operatively by co-registering the CT scan with the implanted assembly to a digital marmoset brain atlas ^47^, from which the schematic probe location was reconstructed (**Fig. 2A3**).

#### Neural Recording and Signal Processing

During each experimental session, which lasted approximately 15 minutes, we used a wireless eCube headstage system (White Matter LLC) to record single-unit activity while animals were freely moving and performing the Mutual Cooperation (MC) task. In total, we recorded 490 neurons from Animal K (14–42 neurons per session; mean ± SEM = 23.3 ± 1.5) and 464 neurons from Animal D (3–43 neurons per session; mean ± SEM = 16.1 ± 2.0). The data logger was attached at the start and removed at the end of each session while the animals were briefly head-fixed, allowing for untethered social interaction during the task. Synchronization between the wireless recordings and the task-control computer was managed by a DAQ board (National Instruments Corp.) connected to the logger’s transceiver. At the start of each recording, the logger sent a pulse that was captured by the NI board and timestamped by the task computer for initial alignment. To correct for any temporal drift, the task computer also sent a continuous 10 kHz signal through the NI board to the logger for auto-correction. Electrophysiological signals were acquired at a 20 kHz sampling rate. The raw data was processed offline: action potentials were first automatically detected and clustered using Kilosort4 ^48^, and the resulting clusters were then manually curated using phy, an open-source Python library, to isolate well-defined single units for all subsequent analyses.

Following spike-sorting, all subsequent analyses were performed on neural data that was first downsampled from 20 kHz to 30 Hz to match the camera’s frame rate. Continuous firing rates were estimated by counting the number of spikes in 33.3 ms (1/30 s) bins and then smoothing the resulting histogram with a *σ* = 50 *ms* Gaussian kernel. This continuous firing rate trace was then z-scored using the mean and standard deviation of that neuron’s activity across the entire session to obtain a normalized firing rate. For the neural DDM, the trial-by-trial slope of firing rate was calculated by fitting a linear regression model to the normalized (z-scored) firing rate trace over the trial’s duration. For the neural GLM (see below), the model was fit directly to the 30 Hz spike count train for each neuron.

### QUANTIFICATION AND STATISTICAL ANALYSIS

#### Computational Modeling of Decision-Making

To prepare the behavioral data for modeling, we first defined trial epochs and key predictor variables. Continuous behavioral data were segmented into discrete trials centered on pull actions. The start of each trial was marked by the onset of the actor’s face linear speed (LS), with response time (RT) defined as the duration until the subsequent pull. Specifically, for each inter-pull interval (defined as the duration between two consecutive pulls by the actor), we computed the median and median absolute deviation (MAD) of LS within that interval, and defined trial onset as the first time point at which LS exceeded the interval-specific median by 3 × MAD. The justification for this marker is detailed in the supplementary materials (**Supp. Fig S1**). To create a holistic measure of the partner’s actions, we performed Principal Component Analysis (PCA) on their eight behavioral variables. These variables were then used to fit a hierarchical Bayesian drift-diffusion model (DDM) to formalize the evidence accumulation process. The DDM decomposes behavior into distinct cognitive parameters: the drift rate ( *v* ), representing the speed of evidence accumulation; the starting point (*z*), representing an initial bias; and the non-decision time (*t_ND_*). To explicitly test our hypothesis about how social gaze mediates the integration of partner information, we formulated and compared four competing models (**Supp. Fig S3A**):

- **M1 (Gaze-gated model):** A model in which partner information influences the drift rate only during periods of social gaze. Periods of social gaze were defined as the time points in the functional window when social gaze probability was greater than zero.

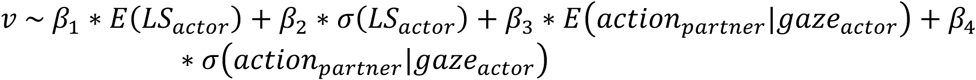
- **M2 (No gaze modulation):** A model where partner information is integrated regardless of gaze.

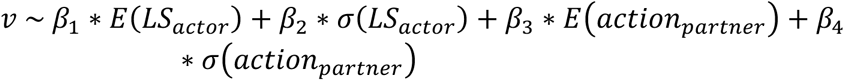
- **M3 (Additive gaze effect):** A model where social gaze (measured as social gaze accumulation, *gaze_auc_*) has a separate, additive effect on the drift rate.

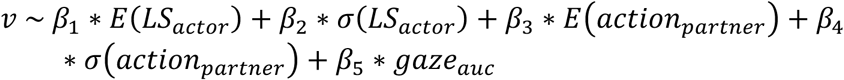
- **M4 (Multiplicative gaze effect):** A model where social gaze multiplicatively scales the influence of partner information.

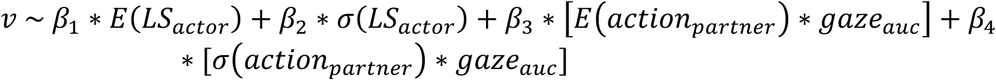
- **M1+ (Gaze-gated model with history effect):** An extension of M1 where the starting point (*z*) is modulated by the outcome of the previous trial, which is 1 if the previous trial is successful, and 0 if it failed. The drift rate (*v*) is formulated identically to M1.

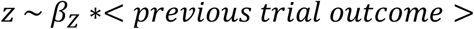
- **M5 (Gaze-gated model with single behavioral variable):** A control model with the same gaze-gated structure as M1, but using the partner’s linear speed (*LS_partner_*) instead of the holistic action variable (PC1, *action_partner_*).

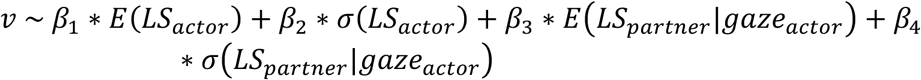

To link the cognitive model to neural activity, we constructed a **neural DDM**. This model used trial-by-trial neural features to predict the DDM parameters. The drift rate (*v*) was predicted by the slope of a neuron’s firing rate, while the starting point (*z*) was predicted by the neuron’s mean baseline firing rate. This baseline activity was calculated during the non-decision time period (*t_ND_*), using the duration estimated for each trial from the fitted parameters of model M1. The specific relationships were:

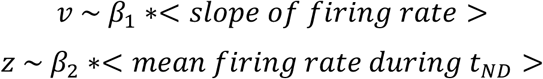

All DDM models were fit using the HDDM (Hierarchical Drift-Diffusion Model) Python package ^21^, which uses Markov Chain Monte Carlo (MCMC) sampling to estimate posterior distributions of the model parameters. For each model, we generated 2000 samples from the posterior distribution following a burn-in period of 1000 iterations, which were discarded. We confirmed model convergence by visually inspecting the MCMC chains. The final coefficient values reported are the mean of the posterior distributions from the collected samples.

#### Social Evidence Estimation

We defined the social evidence for a given trial by subtracting the standard deviation of the partner’s action (PC1) during periods of social gaze from the maximum value observed across all trials, a linear transformation where lower variability (less noise) corresponds to stronger evidence:

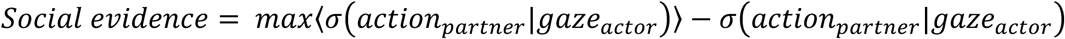

#### Statistical Modeling of Behavior and Neural Tuning

##### Behavioral GLM for Trial Start Identification

To objectively identify the behavioral variable that most reliably predicted the initiation of a pull action, we used a Generalized Linear Model (GLM) with a logistic link function, fit to the eight continuous behavioral variables prepared at a 30 Hz sampling rate. The model was fit separately for each behavioral session. Within each session, the data was split into an 80% training set and a 20% testing set. The model was trained to predict the probability of a pull event from the eight continuous behavioral variables. To capture temporal dynamics, each variable trace was convolved with a basis set of 10 temporal kernels spanning the 4-second window preceding each pull action (−4s to 0s). To determine the importance of each variable, we used a leave-one-out (LOO) procedure. The variable importance was quantified as the drop in the Area Under the Receiver Operating Characteristic curve (AUROC), calculated on the 20% testing set, when that variable was removed from the model. This entire procedure was repeated for 1000 bootstrap iterations to assess the statistical significance of the drop in AUROC for each variable.

##### Neural GLM for Single-Neuron Tuning Properties

We used a Poisson GLM to characterize the tuning of individual dmPFC neurons by fitting the model to the 30 Hz downsampled spike count train for each neuron. The model was fit separately for each neuron using an 80/20 train/test split. The model predicted the number of spikes in each 30 Hz time bin based on four main predictors: the actor’s pull action, social gaze, the partner’s action (PC1), and reward delivery. The reward delivery variable was included as a control to ensure that observed neural activity was not confounded by the motor action of drinking. Each of the four predictors was convolved with a basis set of 10 temporal kernels spanning an 8-second window around each event (−4s to +4s), creating a total of 40 weighted features for the full model.

To determine a neuron’s overall selectivity, we performed an LOO comparison where all 10 kernels for a given variable were excluded. A neuron was considered significantly tuned if the AUROC of the full model on the testing set was significantly better than that of the reduced model. Furthermore, to distinguish between predictive and reactive activity, we assessed them separately based on their temporal relationship to the event under consideration. This was achieved by splitting the 10 temporal kernels into a predictive set (spanning −4s to 0s pre-event) and a reactive set (spanning 0s to +4s post-event). For instance, to test for predictive tuning, we compared the full model to a reduced model where only the 5 predictive kernels for a given variable were excluded. A neuron was classified as having predictive tuning if the full model’s AUROC on the testing set was significantly better than that of this specific reduced model.

#### Population Dynamics

To examine the collective dynamics of the dmPFC population, we carried out single-trial population analyses based on existing methods ^22^. This analysis was restricted to sessions with more than 5 simultaneously recorded neurons (up to 40 neurons per session; 19 sessions in Animal K, 26 sessions in Animal D). For each of these sessions, the normalized (z-scored) firing rates of the population were arranged into an activity matrix. We then performed PCA on each session’s data separately and focused on the neural state space defined by the first three principal components (PCs).

Before calculating geometric measures, each 3D trajectory was preprocessed. First, because trials had different durations, each trajectory was normalized by being reparameterized by its arc length. This process resamples the trajectory so that each path consists of a uniform 100 time points, making them comparable. Second, these reparameterized trajectories were smoothed using a rolling Gaussian window (*σ* = 3) to isolate the large-scale geometry of the path from small-scale neural jitter.

On these preprocessed trajectories, we calculated the following two local geometric measures (**Fig. 4A**):

- **Path Direction (Curvature, *κ*)**: Curvature measures how quickly the trajectory deviates from a straight line. It is formally defined as the magnitude of the rate of change of the curve’s unit tangent vector *T(s)* with respect to arc length *s*.

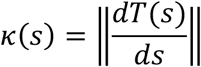
- **Path Variability (Tortuosity, *τ*):** Tortuosity captures the "wiggleness" or complexity of the trajectory. We defined it as the squared ratio of the rate of change of curvature with respect to arc length (*dκ*/*ds*) to the curvature itself.

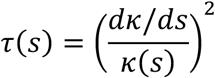

These continuous measures were estimated numerically from the discrete, preprocessed trajectories. Derivatives were approximated using a rolling window of 5 time points, and the term *dκ*/*ds* was calculated using the gradient of the curvature trace. Both the final curvature and tortuosity traces were themselves smoothed with a Gaussian kernel (*σ* = 3) for final analysis.

#### Statistical Analysis

All statistical analyses were performed using custom scripts written in Python with the SciPy and statsmodels libraries. All tests were two-sided, and results were considered significant at p < 0.05. Shaded regions in plots represent the standard error of the mean (SEM).

- **Comparison of Proportions:** A Fisher’s exact test was used to assess the enrichment of predictive tuning in triple-encoding neurons (Fig. 2F). A Chi-squared test was used for other comparisons of categorical data (Fig. 2C, 3C, 3E).
- **Paired Comparisons:** To compare the presence of significant slope encoding on successful versus failed trials within the same neuronal population (Fig. 3D), we used the McNemar test for paired nominal data.
- **One-Sample Tests:** Distributions of coefficients and correlation values were tested for significance against a median of zero using the non-parametric Wilcoxon signed-rank test (Fig. 1H, 3A-B, 3F-G, 4C-D).
- **Time-Series Analysis:** To assess the significance of the group-average time course of path variability (Fig. 4E), we used a permutation test with cluster-based correction. The null hypothesis data was generated by creating one "shuffled" trace for each session. To do this, we first took the real firing rate trace for each neuron in each trial and performed a circular shuffle. The PCA and path variability calculations were then re-run on this shuffled single-trial data to generate the final shuffled trace for each session. The group of real session traces was then compared to the group of shuffled traces using a paired t-test at each time point, with cluster-based correction to control for multiple comparisons.
- **Confirmatory Analyses:** To ensure that results were not confounded by the use of different partners across sessions, all primary statistical analyses were confirmed using linear mixed-effects models with partner identity included as a random effect. These confirmatory models yielded the same statistical conclusions, and therefore, we kept the original results.
- **Successful and failed pull comparison:** Since the successful and failed pulls had different numbers of occurrences, to match the statistical power, we used all failed pull trials and randomly picked the same number of successful pull trials for each session, and performed pair-wise comparisons.

## Acknowledgments

This work was supported by the ‘Biological Sciences Training Program (BSTP)’ training grant (T32 MH014276 to W.S.), the National Institute of Mental Health (R21 MH126072, RF1 MH138396 to S.W.C.C., A.S.N., M.P.J.), the Simons Foundation Autism Research Initiative (SFARI 875855 to S.W.C.C., A.S.N., M.P.J.), Wu Tsai Institute at Yale University (to S.W.C.C., A.S.N., M.P.J., W.S.), a National Eye Institute core grant for vision research (P30 EY026878 to Yale University) and the National Science Foundation Graduate Research Fellowship (DGE2139841 to O.C.M.). We thank the veterinary and husbandry staff at Yale for their excellent animal care, and Anthony DeSimone from the Yale Machine Shop for advice on the neural recording system design.

## Author contributions

Conceptualization: WS, MPJ, SWCC, ASN; Data Collection: WS, OCM; Data Analysis: WS; Computational Modeling: WS; Supervision: MPJ, SWCC, ASN; Writing: WS, MPJ, SWCC, ASN

## Supplemental Figures and Movies

**Figure S1.**
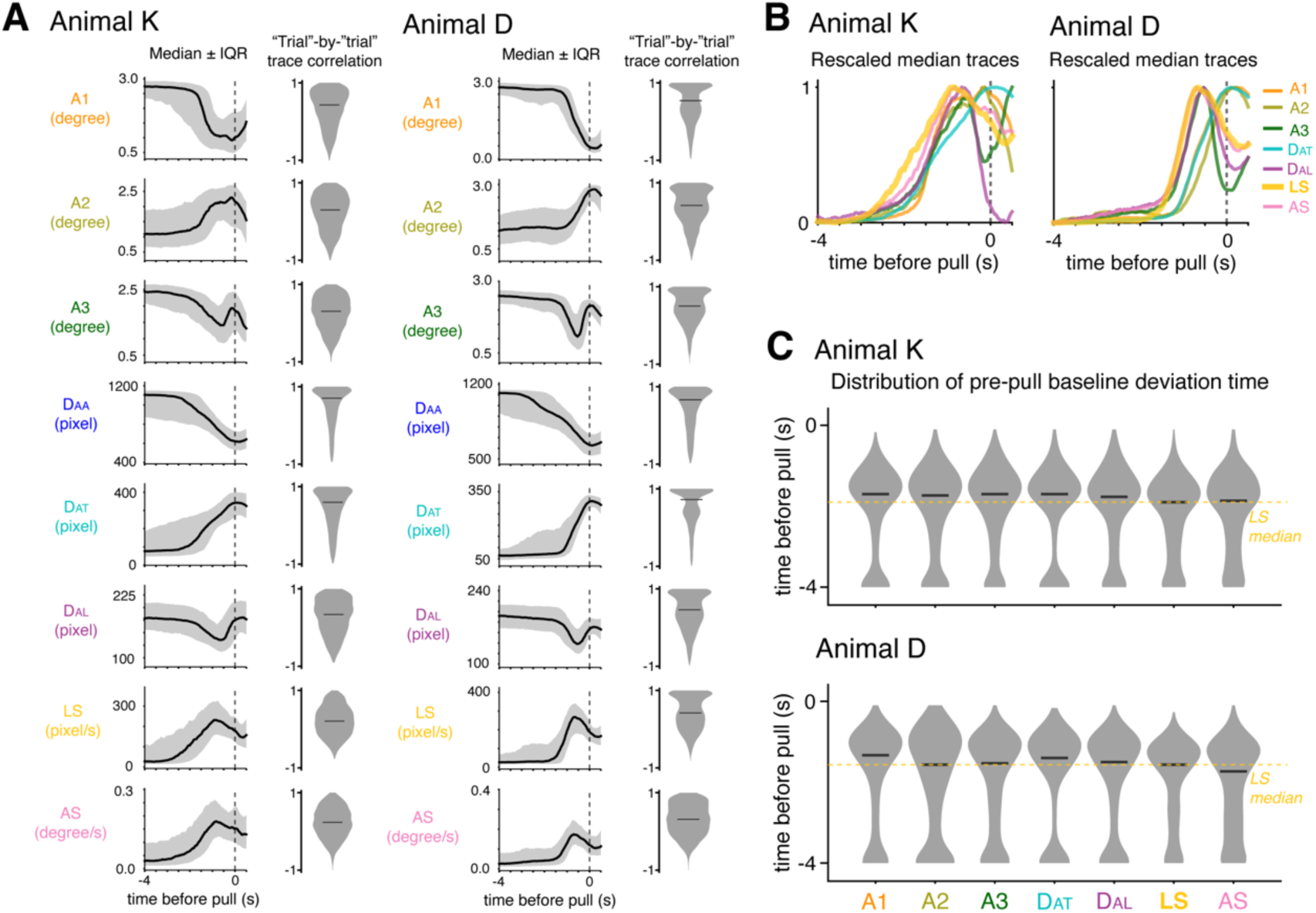
Characterization of continuous behavioral variables preceding pull actions; related to Figure 1. (**A**) Median ± interquartile range time courses aligned to pull onset (left), and the distribution of trial-pair trace correlations computed over the pre-pull window (right). Here, a “trial” refers to the 4-s functional window preceding each pull. (**B**) Rescaled median traces. Median traces of A1, A3, D_AA_ and D_AL_ were sign-flipped, and all traces were scaled to the 0 to 1 range. The distance between animals (D_AA_) is not shown because it depends on both animals’ actions rather than the actor alone. (**C**) Distribution of pre-pull deviation times. For each variable and “trial”, deviation time was defined as the first time point at which the signal exceeded the trial-specific median by three times the median absolute deviation (MAD). The dashed line indicates the median deviation time of linear speed (LS).

**Figure S2.**
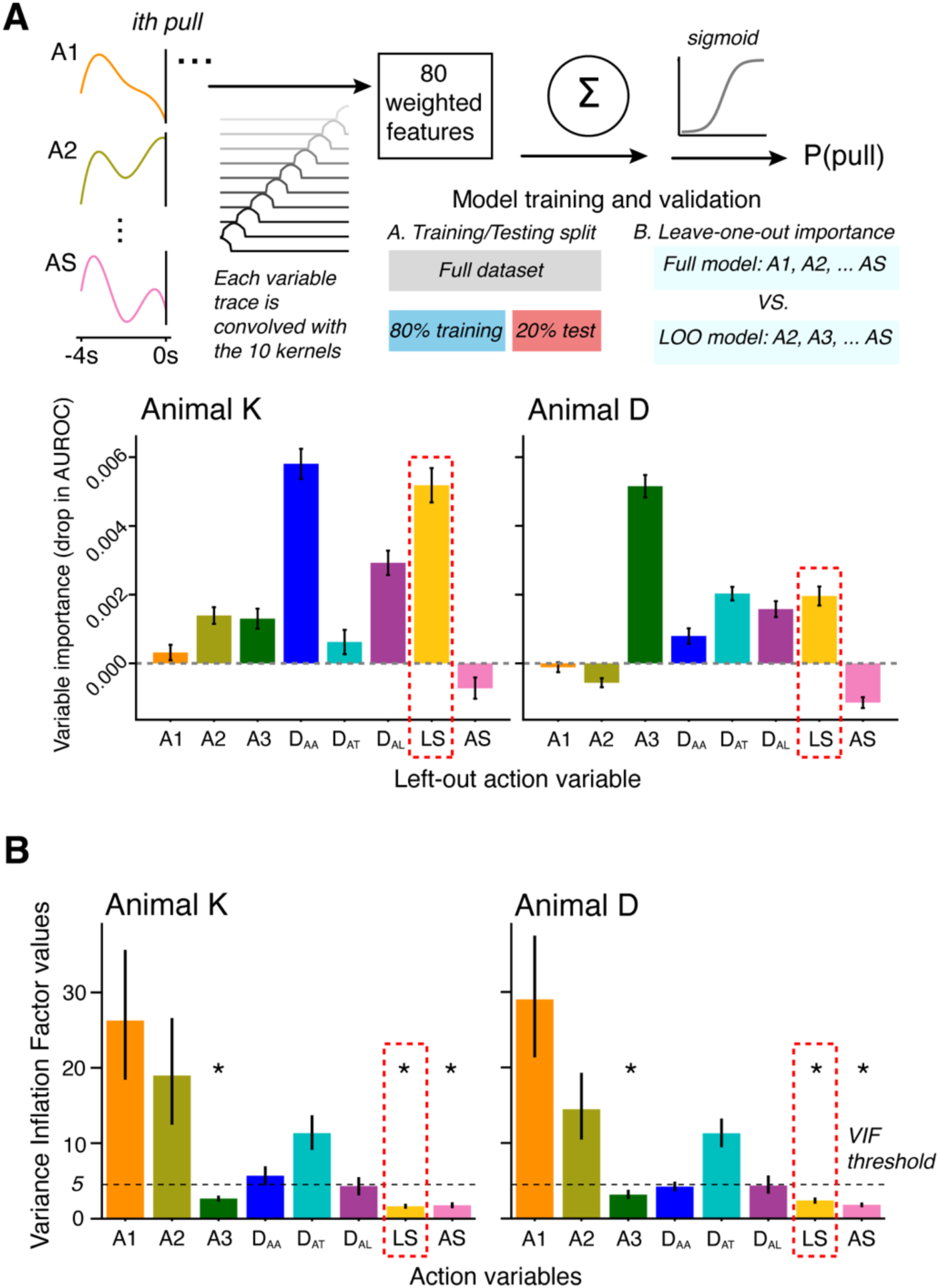
Identifying behavioral variables defining trial start; related to Figure 1. Trial start is best defined by the linear speed (LS) variable. (**A**) Generalized linear model (GLM) pipeline to identify which behavioral variables predict pull actions. (**B**) Collinearity analysis of continuous variables, quantified with variance inflation factors (VIF). Wilcoxon signed-rank test for panel B (* p < 0.05)

**Figure S3.**
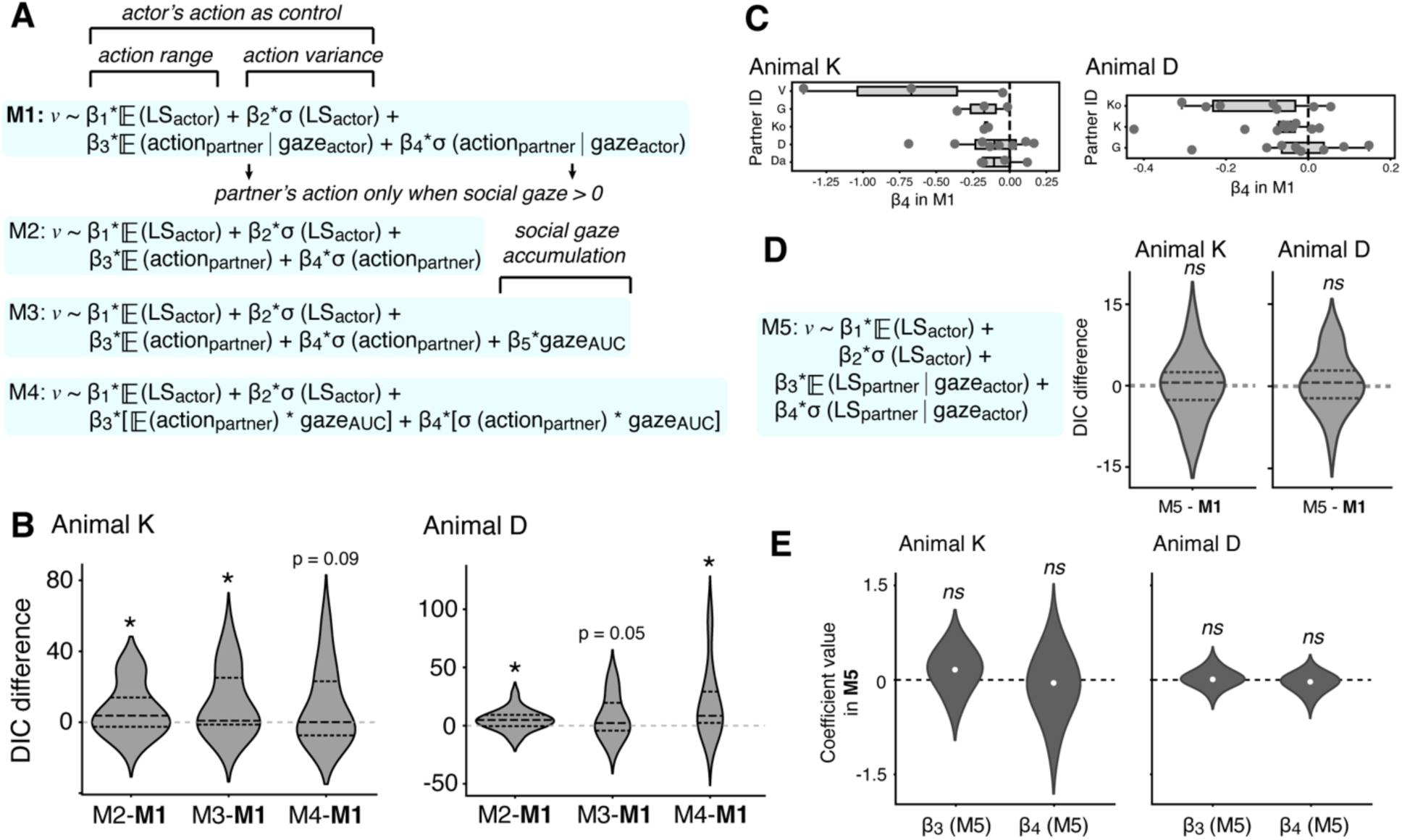
Control drift diffusion models; related to Figure 1. (**A**) Four models testing effects of partner actions and gaze (M1: main model; M2-4: control models). (**B**) Model comparison using Deviance Information Criterion (DIC). (**C**) Partner-wise distribution of the social evidence accumulation coefficient (β4) of the main model M1. (**D**) Comparison of DIC values between the main model (M1, partner PC1 with gaze filter) and control model (M5, partner linear speed, LS, with gaze filter). (**E**) Coefficients from M5, showing non-significant effects. Wilcoxon signed-rank test for panels B-D (ns: no significance, * p < 0.05).

**Figure S4.**
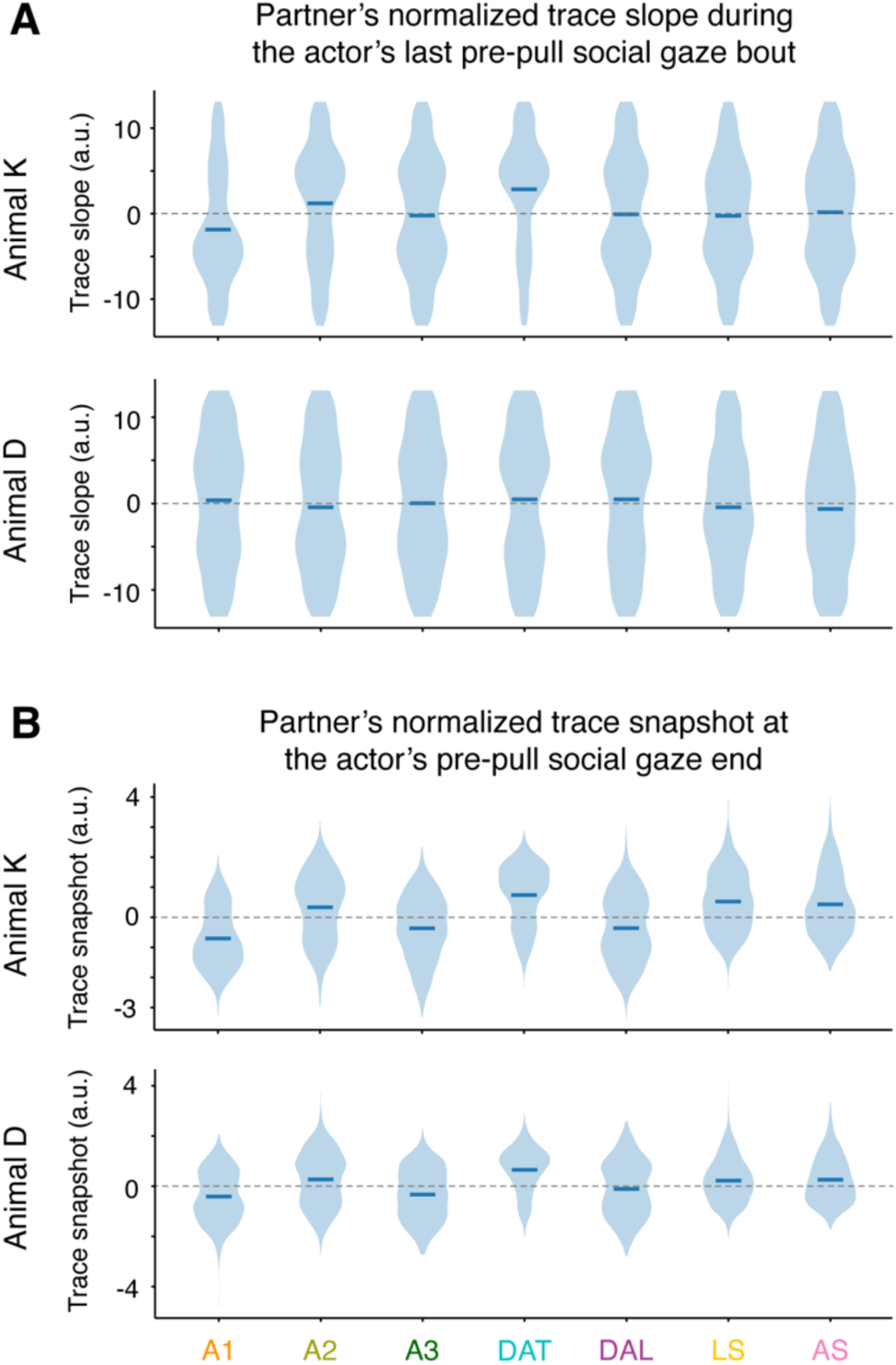
Lack of monotonic trends or fixed partner states during pre-pull social gaze; related to Figure 1. (**A**) Distributions of slopes of partner behavioral variables during the actor’s last pre-pull social gaze bout. For each trial, the slope of each partner behavioral variable was computed during the final social gaze bout occurring prior to pull onset, after within-trial z-scoring. Animal–animal distance (D_AA_) was excluded from this analysis because it reflects a mixture of actor and partner positions and is therefore not purely partner-specific. Across variables, slope distributions are centered near zero, indicating no consistent monotonic trend in partner behavior during social gaze. (**B**) Distributions of end-of-gaze snapshot values for the same behavioral variables. Snapshot values were defined as the mean of the final five video frames (166 ms at 30 Hz) of the last pre-pull social gaze bout after within-trial z-scoring. Snapshot distributions do not cluster at a consistent level across trials, indicating the absence of a fixed partner state associated with gaze termination.

**Figure S5.**
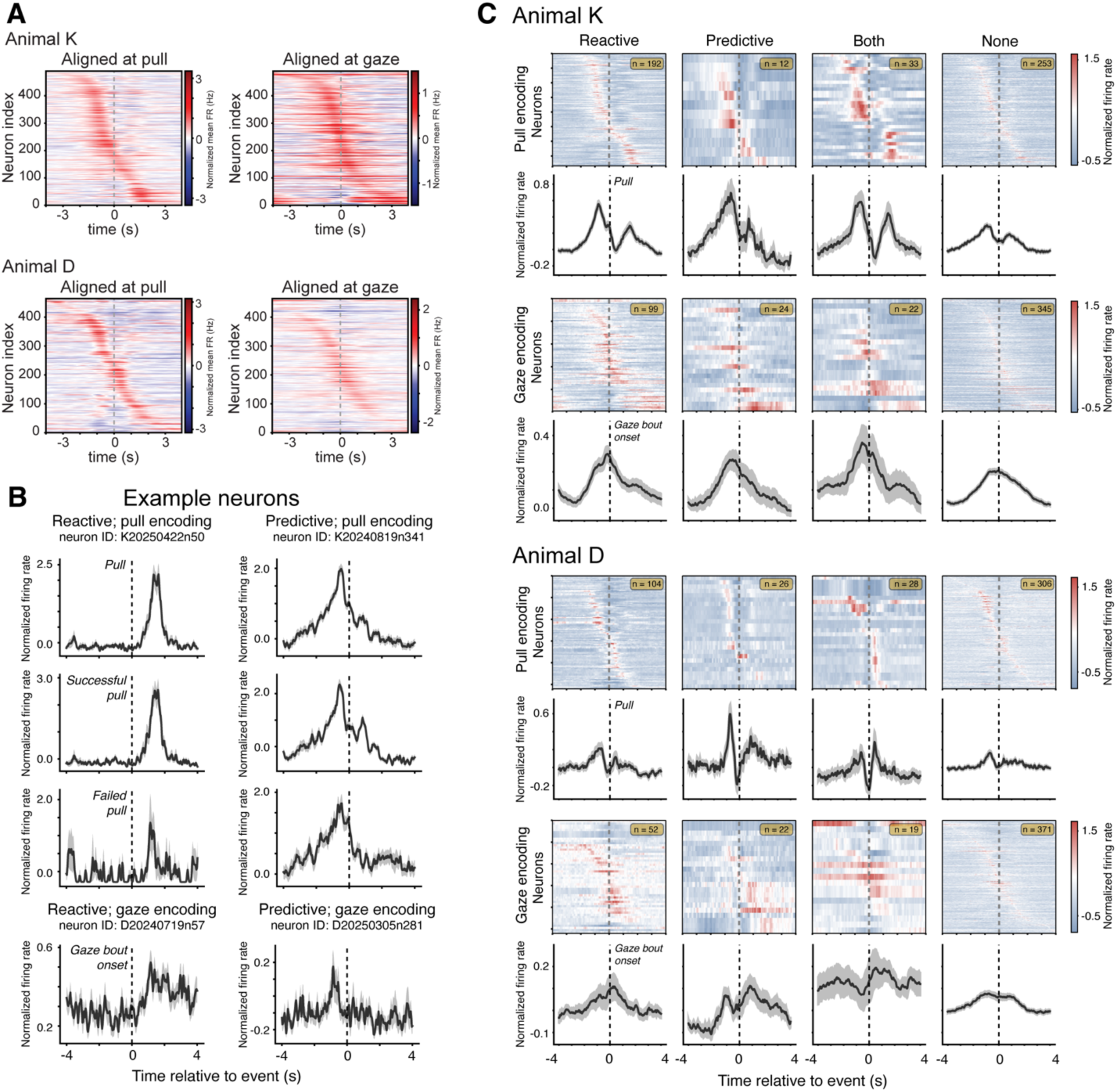
Heatmaps and normalized firing rate traces of dmPFC neural activity; related to Figure 2. (**A**) Heatmaps of normalized (z-scored) firing rates aligned to pull and gaze actions. (**B**) Mean ± SEM traces of normalized (z-scored) firing rates from representative single neurons classified as reactive or predictive for pull or gaze encoding. (**C**) Heatmaps (top) and mean ± SEM traces (bottom) of normalized (z-scored) firing rates grouped by encoding types.

**Figure S6.**
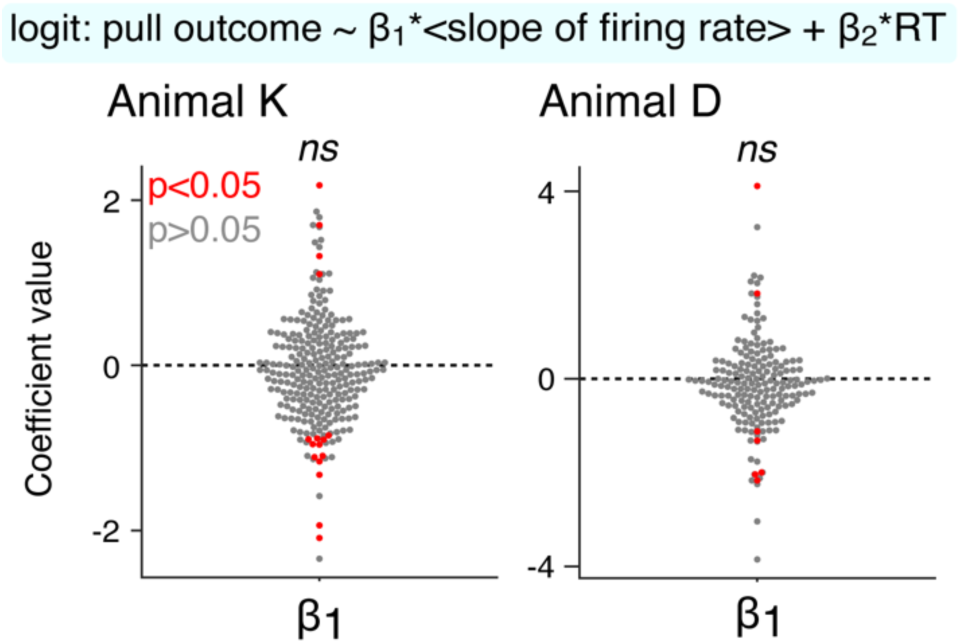
Outcome prediction by ramping slopes after controlling for response time; related to Figure 3. Firing-rate ramp slope coefficients do not predict pull outcome (successful vs. failed) at the population level once response time (RT) is included as a covariate. Each dot represents one neuron–session pair; red dots indicate p < 0.05; dashed lines indicate zero.

**Figure S7.**
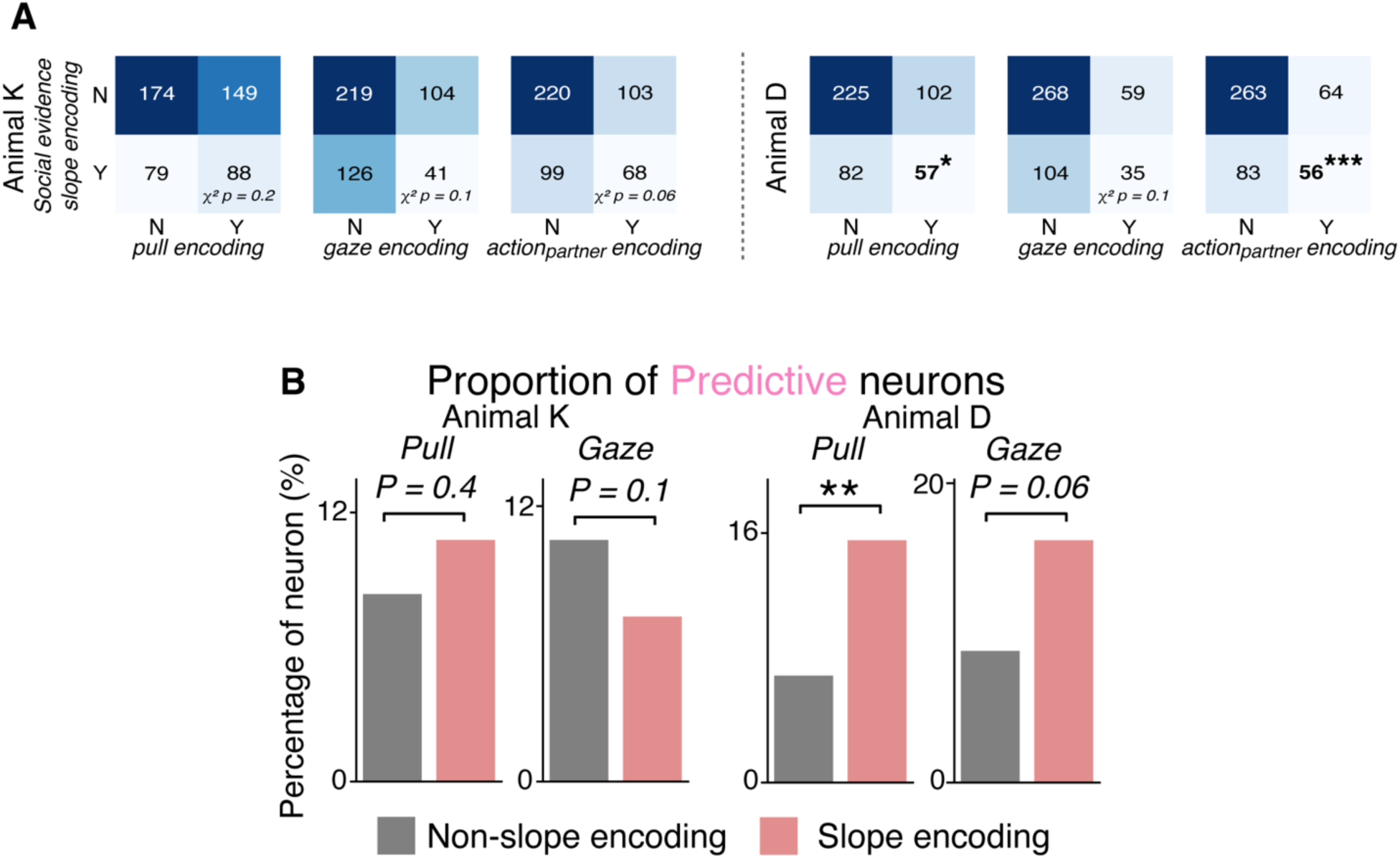
Predictive encoding in slope neurons; related to Figure 3. Slope-encoding neurons are enriched for predictive coding of partner actions. (**A**) Slope-encoding neurons show mixed selectivity for pull, gaze, and partner action variables. (**B**) Proportion of predictive vs reactive encoding among slope and non-slope neurons for pull and gaze. Animal D showed a significant enrichment of predictive coding among slope neurons. Chi-squared test for panel A, Fisher’s exact test for panel B (** p < 0.01).

**Figure S8.**
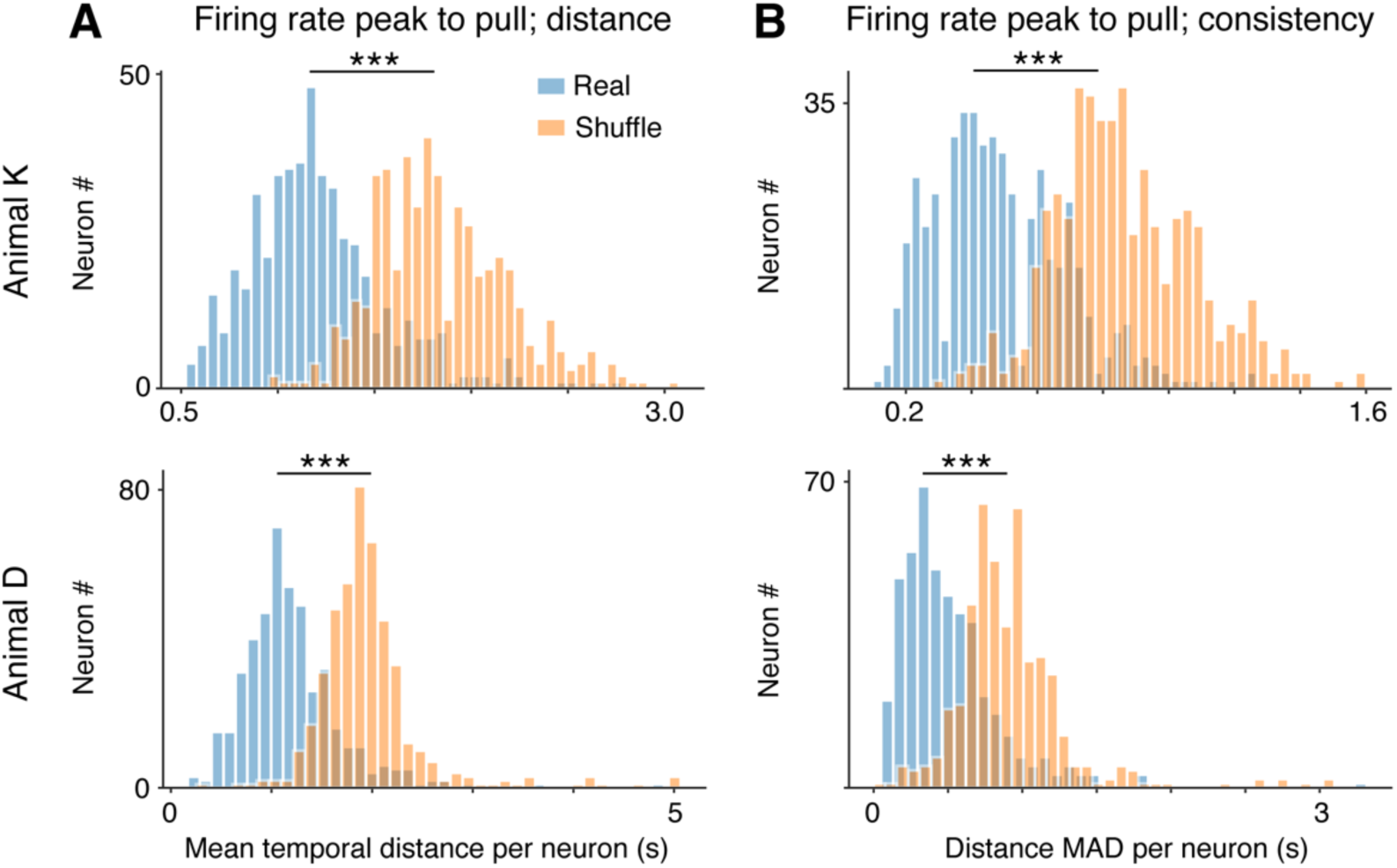
dmPFC firing rate peaks are temporally aligned to the pull action and stable across trials; related to Figure 3. (**A**) Distribution of the mean distance from firing rate peak to pull onset for each neuron, shown separately for the two animals (top: Animal K; bottom: Animal D). Real data (blue) are compared to a control in which the peak time (τ) is randomly assigned within each trial (shuffle; orange). Across both animals, neuronal firing rate peaks occur significantly closer to the pull action than expected under the shuffle control. (**B**) Distribution of the trial-to-trial variability of firing rate peak timing for each neuron, quantified as the median absolute deviation (MAD) of the distance to pull. Real data (blue) show significantly lower variability than the shuffled control (orange), indicating that peak timing is relatively consistent across trials within individual neurons. Paired Student’s t-test for panel A and B (*** p < 0.001).

**Table S1.** Summary statistics of social gaze sampling during successful cooperative trials.

| Animal ID | Animal K | Animal D |
| --- | --- | --- |
| Gaze bouts number per trial<br>mean [min, max] | 1.193 [1.000, 4.000] | 1.104 [1.000, 5.000] |
| gaze bout duration (s)<br>mean [min, max] | 1.491 [0.033, 4.400] | 1.042 [0.033, 4.900] |
| Total gaze duration per trial (s)<br>mean [min, max] | 1.701 [0.033, 5.800] | 1.157 [0.033, 5.467] |
| First gaze bout start (s; rel. pull)<br>mean [min, max] | -1.959 [-7.000, -0.033] | -1.265 [-7.100, -0.033] |
| Last gaze bout end (s; rel. pull)<br>mean [min, max] | -0.173 [-5.900, -0.033] | -0.068 [-3.567, -0.033] |
| RT/trial duration (s)<br>mean [min, max] | 3.618 [1.500, 7.133] | 3.743 [1.633, 7.467] |
| Gaze fraction per trial (total gaze<br>duration/RT)<br>mean [min, max] | 0.508 [0.021, 1.000] | 0.356 [0.008, 1.000] |

**Movie S1. Example clip of the animal tracking and the multi-channel wireless recording form dmPFC.**

## References

1. Burkart, J.M., and van Schaik, C.P. (2010). Cognitive consequences of cooperative breeding in primates? Anim Cogn 13, 1–19. 10.1007/s10071-009-0263-7.

2. Emery, N.J. (2000). The eyes have it: the neuroethology, function and evolution of social gaze. Neurosci Biobehav R 24, 581–604. Doi 10.1016/S0149-7634(00)00025-7.

3. Frischen, A., Bayliss, A.P., and Tipper, S.P. (2007). Gaze cueing of attention: Visual attention, social cognition, and individual differences. Psychol Bull 133, 694–724. 10.1037/0033-2909.133.4.694.

4. Ratcliff, R., Smith, P.L., Brown, S.D., and McKoon, G. (2016). Diffusion Decision Model: Current Issues anc History. Trends in Cognitive Sciences 20, 260–281. 10.1016/j.tics.2016.01.007.

5. Gold, J.I., and Shadlen, M.N. (2007). The neural basis of decision making. Annual Review of Neuroscience 30, 535–574. 10.1146/annurev.neuro.29.051605.113038.

6. Busemeyer, J.R., Gluth, S., Rieskamp, J., and Turner, B. (2019). Cognitive and Neural Bases of Multi-Attribute, Multi-Alternative, Value-based Decisions. Trends in Cognitive Sciences 23, 251–263. 10.1016/j.tics.2018.12.003.

7. Krajbich, I., Armel, C., and Rangel, A. (2010). Visual fixations and the computation and comparison of value in simple choice. Nature Neuroscience 13, 1292–1298. 10.1038/nn.2635.

8. Krajbich, I., and Rangel, A. (2011). Multialternative drift-diffusion model predicts the relationship between visual fixations and choice in value-based decisions. P Natl Acad Sci USA 108, 13852–13857. 10.1073/pnas.1101328108.

9. Hanks, T.D., and Summerfield, C. (2017). Perceptual Decision Making in Rodents, Monkeys, and Humans. Neuron 93, 15–31. 10.1016/j.neuron.2016.12.003.

10. Kim, J.N., and Shadlen, M.N. (1999). Neural correlates of a decision in the dorsolateral prefrontal cortex of the macaque. Nat Neurosci 2, 176–185. 10.1038/5739.

11. Shadlen, M.N., and Newsome, W.T. (2001). Neural basis of a perceptual decision in the parietal cortex (area LIP) of the rhesus monkey. Journal of Neurophysiology 86, 1916–1936. DOI 10.1152/jn.2001.86.4.1916.

12. Meisner, O.C., Shi, W., Fagan, N.A., Greenwood, J., Jadi, M.P., Nandy, A.S., and Chang, S.W.C. (2024). Development of a Marmoset Apparatus for Automated Pulling to study cooperative behaviors. Elife 13. 10.7554/eLife.97088.

13. Meisner, O.C., Shi, W., Nair, A., Nandy, G., Jadi, M.P., Nandy, A.S., and Chang, S.W.C. (2025). Diverse and flexible strategies enable successful cooperation in marmoset dyads. Curr Biol 35, 4509–4521. 10.1016/j.cub.2025.08.005.

14. Weinreb, C., Kannan, L.T., Newman-Boulle, A., Sainburg, T., Gillis, W.F., Plotnikoff, A., Makowska, S., Pearl, J.E., Osman, M.A.M., Linderman, S.W., and Datta, S.R. (2026). Spontaneous behavior is a succession of self-directed tasks. Neuron 114, 922–937. 10.1016/j.neuron.2025.11.021.

15. Ruff, C.C., and Fehr, E. (2014). The neurobiology of rewards and values in social decision making. Nature Reviews Neuroscience 15, 549–562. 10.1038/nrn3776.

16. Gangopadhyay, P., Chawla, M., Dal Monte, O., and Chang, S.W.C. (2021). Prefrontal-amygdala circuits in social decision-making. Nat Neurosci 24, 5–18. 10.1038/s41593-020-00738-9.

17. Isoda, M. (2021). The Role of the Medial Prefrontal Cortex in Moderating Neural Representations of Self and Other in Primates. Annual Review of Neuroscience, Vol 44, 2021 44, 295–313. 10.1146/annurev-neuro-101420-011820.

18. Noritake, A., Ninomiya, T., and Isoda, M. (2018). Social reward monitoring and valuation in the macaque brain. Nature Neuroscience 21, 1452-+. 10.1038/s41593-018-0229-7.

19. Karashchuk, P., Rupp, K.L., Dickinson, E.S., Walling-Bell, S., Sanders, E., Azim, E., Brunton, B.W., and Tuthill, J.C. (2021). Anipose: A toolkit for robust markerless 3D pose estimation. Cell Reports 36. ARTN 109730. 10.1016/j.celrep.2021.109730.

20. Lauer, J., Zhou, M., Ye, S.K., Menegas, W., Schneider, S., Nath, T., Rahman, M.M., Di Santo, V., Soberanes, D., Feng, G.P., et al. (2022). Multi-animal pose estimation, identification and tracking with DeepLabCut. Nat Methods 19, 496–504. 10.1038/s41592-022-01443-0.

21. Wiecki, T.V., Sofer, I., and Frank, M.J. (2013). HDDM: Hierarchical Bayesian estimation of the Drift-Diffusion Model in Python. Front Neuroinform 7. 10.3389/fninf.2013.00014.

22. Monsalve-Mercado, M.M., Stine, G.M., Shadlen, M.N., and Miller, K.D. (2025). The geometry of the neural state space of decisions. bioRxiv. 10.1101/2025.01.24.634806.

23. Dal Monte, O., Fan, S.Q., Fagan, N.A., Chu, C.C.J., Zhou, M.B., Putnam, P.T., Nair, A.R., and Chang, S.W.C. (2022). Widespread implementations of interactive social gaze neurons in the primate prefrontal-amygdala networks. Neuron 110, 2183-+. 10.1016/j.neuron.2022.04.013.

24. Fan, S.Q., Dal Monte, O., and Chang, S.W.C. (2021). Levels of naturalism in social neuroscience research. Iscience 24. 10.1016/j.isci.2021.102702.

25. Jiang, M., Gu, L., Ma, M., Li, Q., Kao, J.C., and Hong, W. (2025). Neural basis of cooperative behavior in biological and artificial intelligence systems. Science, eadw8151. 10.1126/science.adw8151.

26. Jiang, M., Wang, M., Shi, Q., Wei, L., Lin, Y., Wu, D., Liu, B., Nie, X., Qiao, H., Xu, L., et al. (2021). Evolution and neural representation of mammalian cooperative behavior. Cell Rep 37, 110029. 10.1016/j.celrep.2021.110029.

27. Franch, M., Yellapantula, S., Parajuli, A., Kharas, N., Wright, A., Aazhang, B., and Dragoi, V. (2024). Visuo-frontal interactions during social learning in freely moving macaques. Nature. 10.1038/s41586-024-07084-x.

28. Jendritza, P., Klein, F.J., and Fries, P. (2023). Multi-area recordings and optogenetics in the awake, behaving marmoset. Nat Commun 14, 577. 10.1038/s41467-023-36217-5.

29. Walker, J.D., Pirschel, F., Sundiang, M., Niekrasz, M., MacLean, J.N., and Hatsopoulos, N.G. (2021). Chronic wireless neural population recordings with common marmosets. Cell Rep 36, 109379. 10.1016/j.celrep.2021.109379.

30. Calhoun, A.J., Pillow, J.W., and Murthy, M. (2019). Unsupervised identification of the internal states that shape natural behavior. Nature Neuroscience 22, 2040-+. 10.1038/s41593-019-0533-x.

31. Coen, P., Xie, M., Clemens, J., and Murthy, M. (2016). Sensorimotor Transformations Underlying Variability in Song Intensity during Courtship. Neuron 89, 629–644. 10.1016/j.neuron.2015.12.035.

32. Coen, P., Clemens, J., Weinstein, A.J., Pacheco, D.A., Deng, Y., and Murthy, M. (2014). Dynamic sensory cues shape song structure in. Nature 507, 233-+. 10.1038/nature13131.

33. Li, J.W., Aoi, M.C., and Miller, C.T. (2024). Representing the dynamics of natural marmoset vocal behaviors in frontal cortex. Neuron 112. 10.1016/j.neuron.2024.08.020.

34. Chen, R., Radkani, S., Valluru, N., Yoo, S.B.M., and Jazayeri, M. (2026). Evidence accumulation from experience and observation in the cingulate cortex. Nature. 10.1038/s41586-025-09885-0.

35. Tump, A.N., Deffner, D., Pleskac, T.J., Romanczuk, P., and RHJ, M.K. (2024). A Cognitive Computational Approach to Social and Collective Decision-Making. Perspect Psychol Sci 19, 538–551. 10.1177/17456916231186964.

36. Purcell, B.A., and Kiani, R. (2016). Neural Mechanisms of Post-error Adjustments of Decision Policy in Parietal Cortex. Neuron 89, 658–671. 10.1016/j.neuron.2015.12.027.

37. Gold, J.I., Law, C.T., Connolly, P., and Bennur, S. (2008). The Relative Influences of Priors and Sensory Evidence on an Oculomotor Decision Variable During Perceptual Learning. Journal of Neurophysiology 100, 2653–2668. 10.1152/jn.90629.2008.

38. Odoemene, O., Pisupati, S., Nguyen, H., and Churchland, A.K. (2018). Visual Evidence Accumulation Guides Decision-Making in Unrestrained Mice. J Neurosci 38, 10143–10155. 10.1523/JNEUROSCI.3478-17.2018.

39. Fan, S.Q., Dal Monte, O., Nair, A.R., Fagan, N.A., and Chang, S.W.C. (2024). Closed-loop microstimulations of the orbitofrontal cortex during real-life gaze interaction enhance dynamic social attention. Neuron 112. 10.1016/j.neuron.2024.05.004.

40. Camerer, C. (2003). Behavioral game theory: experiments in strategic interaction (Russell Sage Foundation; Princeton University Press). 10.1016/j.socec.2003.10.009.

41. Kennedy, D.P., and Adolphs, R. (2012). The social brain in psychiatric and neurological disorders. Trends in Cognitive Sciences 16, 559–572. 10.1016/j.tics.2012.09.006.

42. Zhu, C.X., Dastani, M., and Wang, S.H. (2024). A survey of multi-agent deep reinforcement learning with communication. Auton Agent Multi-Ag 38. 10.1007/s10458-023-09633-6.

43. Mathis, A., Mamidanna, P., Cury, K.M., Abe, T., Murthy, V.N., Mathis, M.W., and Bethge, M. (2018). DeepLabCut: markerless pose estimation of user-defined body parts with deep learning. Nature Neuroscience 21, 1281-+. 10.1038/s41593-018-0209-y.

44. Singh, V.P., Li, J., Dawson, K., Mitchell, J.F., and Miller, C.T. (2025). Active vision in freely moving marmosets using head-mounted eye tracking. Proc Natl Acad Sci U S A 122, e2412954122. 10.1073/pnas.2412954122.

45. Pandey, S., Simhadri, S., and Zhou, Y. (2020). Rapid Head Movements in Common Marmoset Monkeys. Iscience 23. 10.1016/j.isci.2020.100837.

46. Piza, D.B., Corrigan, B.W., Gulli, R.A., Do Carmo, S., Cuello, A.C., Muller, L., and Martinez-Trujillo, J. (2024). Primacy of vision shapes behavioral strategies and neural substrates of spatial navigation in marmoset hippocampus. Nat Commun 15, 4053. 10.1038/s41467-024-48374-2.

47. Woodward, A., Hashikawa, T., Maeda, M., Kaneko, T., Hikishima, K., Iriki, A., Okano, H., and Yamaguchi, Y. (2022). The Brain/MINDS 3D digital marmoset brain atlas (vol 5, 180009, 2018). Sci Data 9. 10.1038/s41597-022-01247-z.

48. Pachitariu, M., Sridhar, S., Pennington, J., and Stringer, C. (2024). Spike sorting with Kilosort4. Nat Methods 21. 10.1038/s41592-024-02232-7.

